# Programmed ribosomal frameshifting regulates tissue-specific pioneer activity

**DOI:** 10.64898/2026.11.13.766495

**Authors:** Eliana F. Torres-Zelada, Hideyuki Komori, Marcel Morgenstern, Benjamin Chadwick, Hsiao-Yun Liu, Elizabeth D. Larson, Ally W.H. Yang, Timothy R. Hughes, Andrew A. Mehle, Joshua J Coon, Christine A. Rushlow, Cheng-Yu Lee, Melissa M. Harrison

## Abstract

Pioneer transcription factors bind nucleosomal DNA and establish accessible chromatin, yet their binding is tissue specific^1–5^. While studies have largely focused on protein-extrinsic features, like chromatin structure and cofactor expression, the mechanisms governing the tissue-specific binding remain poorly understood. We used the *Drosophila* pioneer transcription factor Zelda as a paradigm to investigate the mechanisms controlling tissue-specific binding. Zelda promotes the undifferentiated state in both the embryo and in neural stem cells of the larval brain^6–10^. However, Zelda binds thousands of unique sites in each tissue and, in so doing, defines tissue-specific enhancers^6^. We showed that tissue-specific binding is controlled by the developmentally regulated use of two distinct DNA-binding domains. Our identification of a novel, neural stem cell enriched Zelda proteoform that lacks the embryo-specific C-terminal DNA-binding domain explains the differential DNA-binding domain usage. This proteoform arises through developmentally regulated −1 ribosomal frameshifting, a mechanism common to viruses but not documented for any eukaryotic transcription factors. Together we revealed a new strategy where developmentally programmed ribosomal frameshifting confers tissue-specific pioneer-factor binding and suggest that this mechanism may be more broadly used to diversify the proteome than previously appreciated.

---

Coordinated changes in gene expression drive the development of a complex organism from a single-celled zygote. Transcription factors mediate these changes in cellular identity by binding DNA and regulating specific gene-expression programs. While chromatin is a barrier to DNA binding by many transcription factors, specialized pioneer transcription factors bind nucleosomal DNA, establish regions of open chromatin, and facilitate the binding of additional transcription factors. In so doing, pioneer factors define the *cis*-regulatory regions that control gene expression^11,12^. Because of their ability to restructure the chromatin accessibility landscape, pioneer factors act at the top of gene-regulatory networks to control developmental transitions^13–16^. Although pioneer factors bind nucleosomal DNA, their capacity to bind and open chromatin is not unrestrained. Only a subset of recognition motifs is bound *in vivo*, and binding is tissue specific^1–5^. Therefore, pioneer factors both regulate development and are influenced by developmental context^14,16,17^. Nonetheless, the features that regulate tissue-specific pioneer-factor occupancy remain unclear.

Because of their ability to promote broad changes in gene expression, pioneer factors drive conserved developmental transitions. One of these transitions is the initial stages of development in which the specified germ cells are reprogrammed to totipotency. At this time, the zygotic genome is transcriptionally quiescent, and maternally deposited mRNAs and proteins control the first hours of development. Widespread transcription is initiated only after the cells have become totipotent^18,19^. In all organisms studied to date, pioneer factors promote reprogramming of the zygotic genome and transcriptional activation^16,17,20,21^. The first such pioneer factor identified was Zelda (Zld) in *Drosophila melanogaster*^7–10^. Since its identification, Zld has served as a paradigm for understanding pioneer-factor mediated reprogramming.

In addition to its essential function in embryos, Zld promotes the undifferentiated fate in neural stem cells (neuroblasts) of the larval brain. Neuroblasts divide asymmetrically to both self-renew and generate progeny that rapidly exits the multipotent state and begins to differentiate^22^. Failure to robustly control this transition leads to excess stem-cell divisions and tumors^23,24^. Indeed, overexpression of Zld generates extra neuroblasts and a tumor-like phenotype^6^. Despite the shared role in promoting the undifferentiated state in embryos and neuroblasts, Zld occupies distinct regions of the genome in these two cell populations^6^. In embryos, Zld binding is driven strongly by sequence with 90% of the most highly bound regions containing the canonical CAGGTA motif^7^. By contrast, Zld-bound regions in neuroblasts are only weakly enriched for a degenerate version of this motif^6^. Thus, Zld has tissue-specific genome occupancy, like other pioneer factors^1–5^. However, the mechanisms driving the distinctive binding in embryos and neuroblasts remain unclear.

Protein-extrinsic features, like chromatin structure and cofactor expression, have been thought to be the predominant regulators of tissue-specific pioneer-factor binding^1–5,25–29^. Contrary to expectations, we identified a protein-intrinsic feature that governs the tissue-specific binding: production of a novel, neuroblast-enriched Zld proteoform via developmentally regulated −1 ribosomal frameshifting. The frame-shifted proteoform lacks the canonical C-terminal CAGGTA-binding domain, but retains a second, previously unappreciated DNA-binding domain in Zld required for binding and transcriptional activation in neuroblasts. Our discovery of ribosome frameshifting as a regulator of DNA-binding domain usage reveals a novel mechanism controlling tissue-specific pioneer-factor activity and suggests this process may be used broadly to diversify the proteome across tissues.

## The deeply conserved second zinc finger in Zld has tissue-specific functionality

Eukaryotic transcription factors are commonly comprised of a single, structured DNA-binding domain and extensive intrinsically disordered regions^30^. Both the structured DNA-binding domain and the disordered regions contribute to chromatin binding^31^. Like other factors, Zld contains a structured, C-terminal cluster of four C2H2 zinc fingers (ZnF) that form the DNA-binding domain required for activating embryonic Zld-target genes and is otherwise largely unstructured (Fig. 1a)^32^. Zld also contains two, structured, N-terminal zinc fingers. ZnF2 is the most deeply conserved region of the protein and functions to inhibit Zld-mediated activation of zygotic transcription^33,34^.

**Figure 1:**
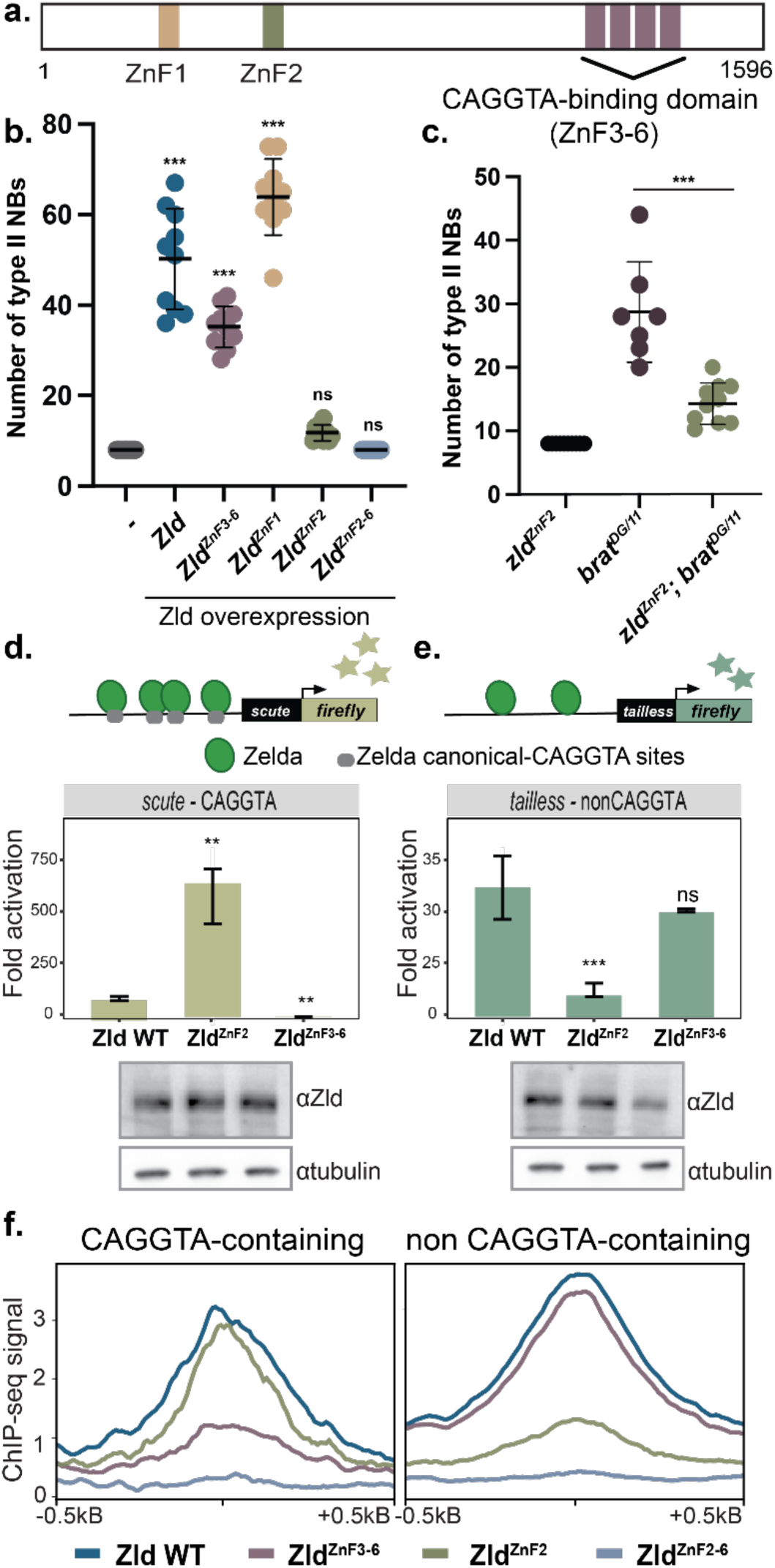
The second zinc finger of Zld regulates tissue-specific activity. **a.** Schematic of Zld including the six zinc fingers. **b.** Number of type II neuroblasts (NBs) per lobe of brains in which the indicated Zld constructs are driven by *Wor-Gal4, tub-Gal80^ts^*. Comparisons are to *Wor-Gal4, tub-Gal80^ts^* alone using an ANOVA. **c.** Number of type II neuroblasts per brain lobe in the genotypes indicated. Males were used in all assays to allow selection of wild-type or hemizygous *zld^ZnF2^* larvae. Genotypes were compared using an unpaired t-test. For each genotype: n = 10 brains. **d.** Fold activation of the *scute* luciferase reporter driven by expression of wild-type (WT), ZnF2-mutant or ZnF3-6 mutant Zld as compared to an empty expression vector. The *scute* promoter contains four CAGGTA elements. **e.** Fold activation of the *tailless* luciferase reporter driven by expression of wild-type (WT), ZnF2-mutant or ZnF3-6 mutant Zld as compared to an empty expression vector. The *tailless* reporter contains a previously identified type II neuroblast-specific enhancer. Comparisons are made to activation by wild-type Zld using a t-test. n = 3 technical replicates. Immunoblots for Zld demonstrate equal protein levels. Tubulin is included as a loading control. **f.** Metaplots of Zld ChIP-seq in S2 cells expressing the Zld proteins indicated below. Signal is shown for binding to regions of closed chromatin containing CAGGTA sites (*left*) or not containing CAGGTA sites (*right*). **, p < 0.01. ***, p < 0.001. ns = not significant.

Zld promotes the undifferentiated fate in both the early embryo and neuroblasts in the larval brain^6,8^. There are exactly eight type II neuroblasts per lobe in the larval brain. Thus, extra type II neuroblasts are indicative of changes to cell fate. Expression of exogenous Zld results in extra type II neuroblasts, and this depends, in part, on ZnF3-6 (Fig. 1b)^6^. As ZnF2 has inhibitory activity in the embryo^33^, we predicted that mutations in this domain would further enhance the ability of Zld to promote extra type II neuroblasts. However, mutations in ZnF2 completely abrogated Zld-mediated type II neuroblast induction (Fig. 1b). This counterintuitive result reflects the endogenous function of ZnF2. Mutations in *brain tumor* (*brat*) cause a tumor-like phenotype that requires Zld^6,24^. Males hemizygous for a mutation in ZnF2 suppressed the extra type II neuroblasts in *brat* hypomorphic mutant brains, phenocopying *zld* null alleles (Fig. 1c). Together, these data reveal that the function of ZnF2 is context-dependent: required to promote the type II neuroblast fate, but inhibiting the role of Zld in embryonic gene expression.

## Underlying DNA sequence and not cofactor expression regulates ZnF2 activity

A potential explanation for the opposing functions of ZnF2 in embryos and type II neuroblasts is the different cellular contexts, including chromatin state and cofactor expression. To test the influence of the cellular environment on ZnF2 function, we exogenously expressed Zld in S2 cell culture and determined the ability of Zld to activate transcription of either an embryonic reporter (*scute* (*sc*)), which contains four canonical CAGGTA motifs, or a type II neuroblast reporter (*tailless* (*tll*)), which lacks these motifs^6,32^. Zld robustly activated transcription of both reporters (Fig. 1d,e). Expression of Zld with mutations in ZnF2 (Zld^ZnF2^) hyperactivated the *scute* reporter, while mutations to ZnF3-6 (Zld^ZnF3–6^) abrogated this activity (Fig. 1d). By contrast, Zld-mediated activation of the *tll* reporter depended on ZnF2, but was independent of ZnF3-6 (Fig. 1e). Because reporter identity, not cell type, dictated the dependence on ZnF2, the activity of ZnF2 is controlled by the underlying DNA-regulatory element.

We used chromatin immunoprecipitation coupled with high-throughput sequencing (ChIP-seq) to determine how mutations to ZnF2 or ZnF3-6 influenced chromatin binding in S2 cells. Zld binds promiscuously to accessible chromatin^26^. Therefore, we focused our initial analysis on regions at which Zld functions as a pioneer factor by binding to closed chromatin and promoting accessibility26. We subdivided these Zld-pioneered sites depending on whether they contained a canonical CAGGTA Zld-binding motif (Fig. 1f). Mutations in the previously defined DNA-binding domain (ZnF3-6) resulted in a loss of chromatin binding at regions containing a CAGGTA motif, but this mutant retained binding at loci lacking the canonical motif (Fig. 1f). By contrast, ZnF2 was dispensable for binding CAGGTA-motif containing regions but required for binding other loci (Fig. 1f). Analysis of all ChIP-seq peaks confirmed that mutations in ZnF3-6 result in a loss of Zld binding at sites enriched for the CAGGTA motif (WT unique, Extended data Fig. 1a.) When ZnF2 was mutated, sites that gained binding were enriched for the CAGGTA motif, consistent with an inhibitory role for ZnF2 in displacing Zld from CAGGTA motifs (ZnF2 unique, Extended data Fig. 1b). As expected, Zld binding was lost at all Zld-pioneered sites when ZnF2-6 were mutated (Fig. 1f). These results suggest that ZnF2 and ZnF3-6 mediate distinct and independent activities that regulate DNA binding, with ZnF3-6 promoting binding to CAGGTA elements and ZnF2 promoting binding to noncanonical elements. The identification of this small, structured domain as influencing sequence-binding preference stands in contrast to the recent focus on the intrinsically disordered regions.

## The DNA-binding activity of Zld is mediated by two distinct, structured domains with unique motif preferences

To determine if ZnF2 directly interacts with DNA, we assessed binding specificity using protein binding microarrays (PBMs) with purified recombinant regions of Zld, either the N-terminus with ZnF2 (Zld N) or the C-terminus with ZnF3–6, (Zld C) (Fig. 2a)^35^. As expected, Zld C preferentially bound the canonical CAGGTA motif (Fig. 2a)^36^. By contrast, Zld N showed the highest affinity for a G-rich variant motif (AAGGGT) (Fig. 2a). To validate binding of Zld N to the newly identified G-rich motif, we performed electrophoretic mobility shift assays (EMSAs) using a CAGGTA-containing probe derived from the *scute* promoter^7,32,37^. Binding specificity was determined by replacing the CAGGTAG site with the AAGGGTT motif. Zld C specifically bound to the CAGGTA-containing probe, while Zld N preferentially bound to the AAGGGT variant (Fig. 2b; Extended data Fig. 2). Neither protein bound to a mutated probe (Fig. 2b). Binding by Zld N to the AAGGGT-containing probe depended on ZnF2, as a ZnF2 mutant exhibited a ∼10-fold decrease in binding (Fig. 2c). These *in vitro* studies demonstrate a direct interaction between ZnF2 and the AAGGGT motif. This is further supported by our S2 ChIP-seq data showing that ZnF2 was required for binding to AAGGGT motifs in closed chromatin, whereas ZnF3-6 were dispensable (Fig. 2d). These differences in binding are functionally relevant, as substitution of the four CAGGTAG sites to AAGGGTT converted the *scute* reporter from depending on ZnF3-6 for activation to depending on ZnF2 (Fig. 2e), essentially making this reporter mimic the type II neuroblast-specific *tll* reporter. Thus, ZnF2 is a previously unidentified DNA-binding domain in the paradigmatic pioneer factor Zld.

**Figure 2:**
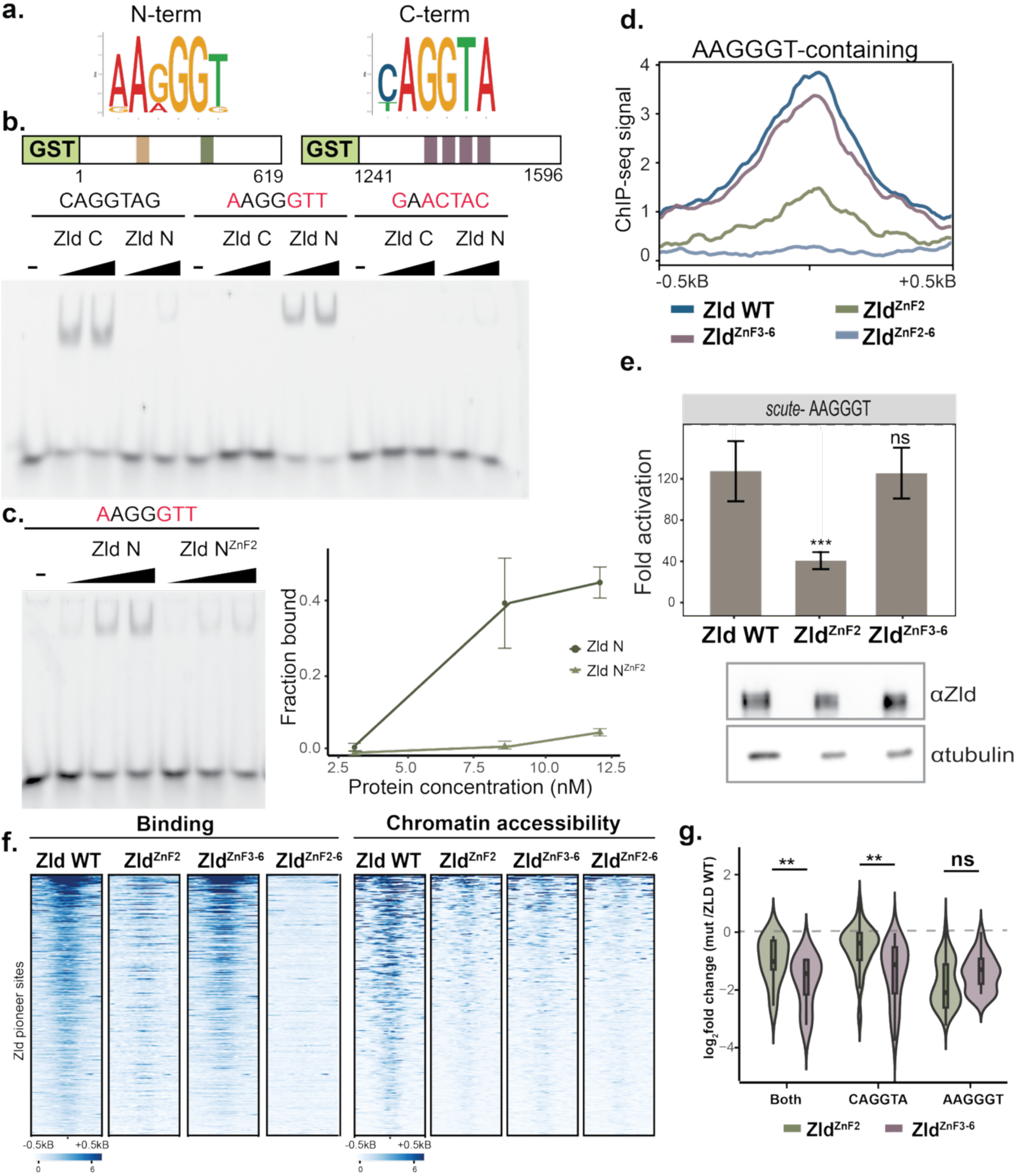
Zld contains two DNA-binding domains with distinct sequence specificities. **a.** Position-weight matrices from protein-binding microarrays based on binding by recombinant Zld constructs. Zld N is an N-terminal construct containing ZnF1 and ZnF2. Zld C is a C-terminal construct contain ZnF3-6. **b.** Schematics of Zld constructs used for EMSA (*top*). Representative EMSA showing binding of Zld N or Zld C to probes containing the motifs indicated above. Red highlights bases that differ from the CAGGTAG motif. **c.** Representative EMSA of binding by Zld N or Zld N with mutations in ZnF2 to an AAGGGT-containing probe (*left*). Fraction of DNA bound as a function of protein concentration for two replicate EMSAs (*right*). Data points are the mean ± standard deviation. **d.** Metaplots of Zld ChIP-seq in S2 cells expressing the Zld proteins indicated below. Signal is shown for binding to regions of closed chromatin containing AAGGGT motifs. **e.** Fold activation of a *scute* luciferase reporter in which all four CAGGTAG motifs are replaced by AAGGGTT. Activation is driven by expression of wild-type (WT), ZnF2-mutant or ZnF3-6 mutant Zld. Comparisons are made to wild-type Zld using a t-test. n = 3 technical replicates. Immunoblots for Zld demonstrate equal protein levels. Tubulin is included as a loading control. **f.** Heatmaps of z-score normalized Zld ChIP-seq (*left*) and ATAC-seq (*right*) in S2 cells expressing the Zld proteins indicated above. Heatmaps are centered on peak summits (± 500 bp). **g.** Violin plots of log_2_ fold change in ATAC-seq signal (Zld^ZnF2^ or Zld^ZnF3–6^ vs wild-type Zld) at all Zld-bound pioneered sites (Both) or subdivided into regions that contain either CAGGTA motifs or AAGGGT motifs (as indicated below). t-tests were used for comparisons between genotypes. **, p < 0.01. ***, p < 0.001. ns = not significant.

Binding and opening closed chromatin are defining features of pioneer factors controlled, in part, by their DNA-binding domains. We therefore investigated how ZnF2 and ZnF3-6 each contribute to the capacity of Zld to promote chromatin accessibility. By coupling ChIP-seq with analysis of assay for transposase accessible chromatin (ATAC)-seq, we previously showed that Zld promotes chromatin accessibility at a subset of bound regions in S2 cells^26^. We used this same approach to show that mutation of ZnF2 abrogated binding and the ability of Zld to generate chromatin accessibility at this subset of bound regions (Fig. 2f). Zld^ZnF3–6^ largely retained binding. However, binding by Zld^ZnF3–6^ did not promote chromatin accessibility (Fig. 2f). When analyzed genome wide, mutations in ZnF3-6 had a significantly more pronounced effect on chromatin accessibility than mutation of ZnF2, despite the broader requirement for ZnF2 for DNA binding (Fig. 2g). Thus, ZnF2 is required for DNA binding but ZnF3-6 is needed for pioneering activity.

## ZnF2 is a neuroblast-specific DNA-binding domain

Our identification of ZnF2 as a newly identified DNA-binding domain specifically required in type II neuroblasts suggested that differential usage of the two DNA-binding domains in Zld could drive the tissue-specific chromatin occupancy. We therefore performed CUT&RUN to test the role of ZnF2 in regulating Zld binding in type II neuroblasts. Binding sites for wild-type or mutant versions of Zld were identified in *brat* null-mutant brains enriched for thousands of type II neuroblasts^6^. ZnF2, but not ZnF3-6 was broadly required for Zld occupancy in type II neuroblasts, including at the type II neuroblast-specific *tll* enhancer (Fig. 3a,b). Notably, ZnF2-dependent sites were enriched for the AAGGGT motif (Fig. 3c). Exogenously expressed wild-type Zld binds to largely the same loci as endogenous Zld (Fig. 3a, Extended data Fig. 3). As predicted for a pioneer factor, exogenous expression resulted in additional binding sites, but comparing between exogenously expressed wild-type and mutant protein allowed us to control for these novel binding sites (Fig. 3a, Extended data Fig. 3d). Importantly, the HA-epitope used for CUT&RUN did not affect Zld binding or activity (Extended data Fig. 3).

**Figure 3:**
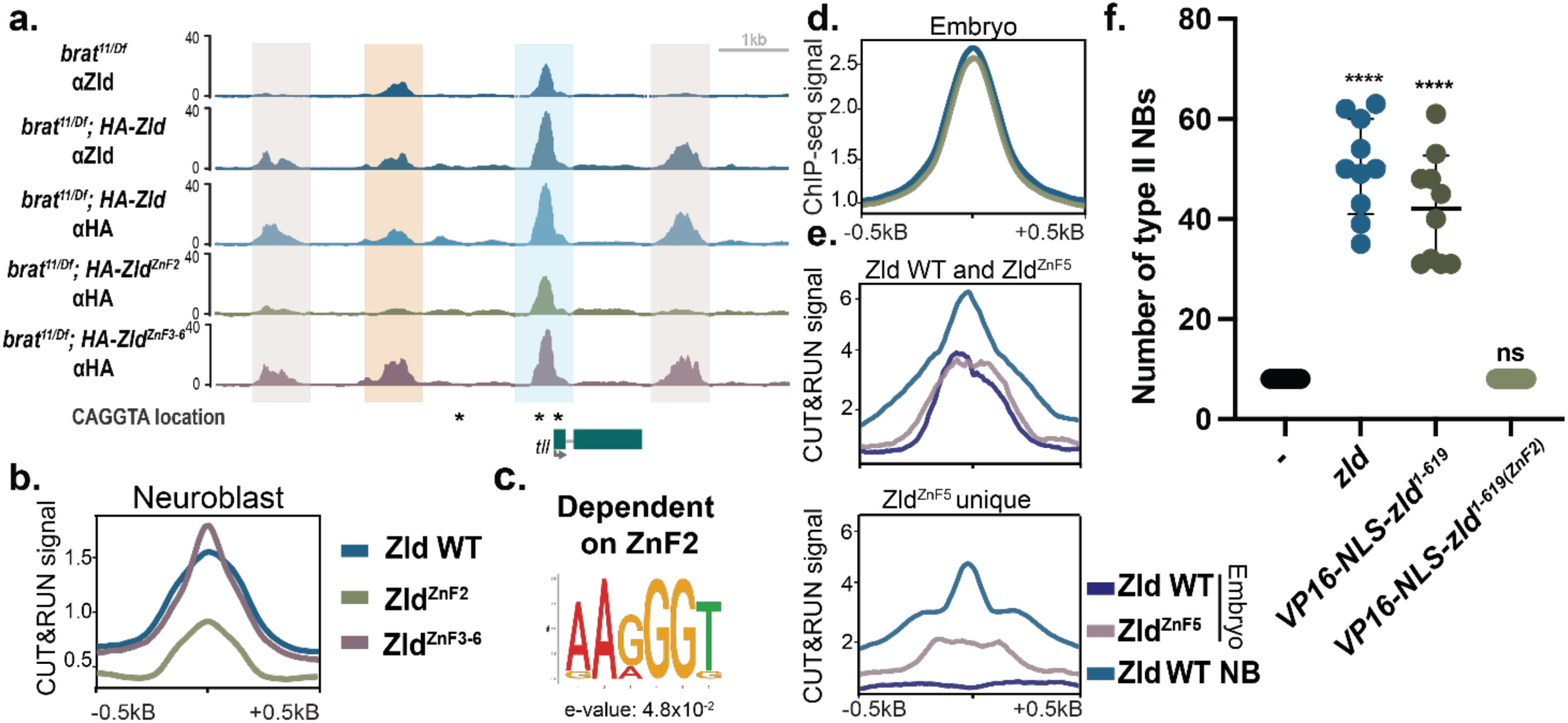
ZnF2 confers tissue-specific chromatin binding that is necessary and sufficient for promoting neuroblast fate. **a.** Genome browser tracks over the *tll* locus of CUT&RUN against either HA or Zld. The type II neuroblast-specific enhancer is highlighted in orange, the promoter in blue, and regions that are bound by Zld only upon exogenous expression in gray. **b.** Metaplots of average z-score normalized anti-HA CUT&RUN from brains expressing the indicated HA-tagged Zld proteins. **c.** Position weight matrix of the AAGGGT motif enriched in regions that depend on ZnF2 in type II neuroblasts (determined by MEME-suite). **d.** Metaplots of spike-in normalized ChIP-seq for Zld from either wild-type or *zld^ZnF2^* embryos **e.** Metaplots of z-score normalized CUT&RUN for Zld from wild-type or *zld^ZnF5^* embryos (from Fallacaro et al. bioRxiv 2024) or wild-type type II neuroblasts. Regions bound by both wild-type Zld and Zld^ZnF5^ (*above*). Regions unique to Zld^ZnF5^ (*below*). **f.** Number of type II neuroblasts per brain lobe in which the indicated VP16-NLS Zld constructs are exogenously expressed. Comparisons are to *Wor-Gal4, tub-Gal80^ts^* alone using an ANOVA. ****, p < 0.0001. ns = not significant.

In contrast to the type II neuroblasts, the CAGGTA motif is a strong predictor of Zld occupancy in embryos^6,7,38^. This suggests AAGGGT recognition by ZnF2 does not drive Zld binding at this early stage of development, and its activity might impede the recruitment of Zld to embryo-specific *cis*-regulatory elements. To test this, we performed spike-in normalized ChIP-seq for Zld in wild-type embryos (*w*^1118^) or embryos laid by mothers homozygous for *zld^ZnF2^* ^33^. Mutation of ZnF2 did not significantly change the genome occupancy of Zld in the embryo (Fig. 3d, Extended data Fig. 4a), indicating that ZnF2 is selectively required for binding in type II neuroblasts but dispensable for its genome-wide binding in the embryo. Importantly, Zld and Zld^ZnF2^ were expressed at equivalent levels (Extended data Fig. 4b). While genes upregulated in *zld^ZnF2^* embryos might be expected to show higher Zld occupancy when ZnF2 was mutated, this was not the case (Extended data Fig. 4c)^33^. We then determined if ZnF2 promoted chromatin occupancy in the embryo as it did in type II neuroblasts by reanalyzing CUT&RUN data from embryos in which ZnF5 was mutated to abrogate binding to the canonical CAGGTA motif^39^. Indeed, regions bound by Zld^ZnF5^ in the embryo (both unique and shared with wild type) were strongly bound by Zld in type II neuroblasts and depended on ZnF2 for this binding (Fig. 3e, Extended data Fig. 4d). Thus, loci specifically bound in the ZnF5 mutant embryos likely indicate new binding events driven by ZnF2.

Our data support a model in which competition between two, distinct DNA-binding domains (ZnF2 and ZnF3-6) results in tissue-specific chromatin occupancy. In the embryo, ZnF3-6 dominates, while in the type II neuroblasts ZnF2 is required. We therefore tested if ZnF2 was sufficient to promote Zld occupancy at Zld-responsive loci in type II neuroblasts. We fused the N-terminal region of Zld (aa 1-691), containing ZnF2 but not ZnF3-6, to the potent VP16 transcriptional activation domain and included a nuclear localization signal (NLS) to confer the potential for transcriptional activation (Extended data Fig. 5a)^40,41^. VP16-NLS-Zld^1–619^ activated both the *scute* reporter with AAGGGT motifs and the *tll* reporter in a ZnF2-dependent manner, confirming the ability of this protein to promote transcription (Extended data Fig. 5b). ZnF2 was also sufficient to drive extra type II neuroblasts in the animal. Expression of VP16-NLS-Zld^1–619^ in type II neuroblasts resulted in extra neuroblasts similar to the full-length protein, and this effect was dependent on ZnF2 (Fig. 3f, Extended data Fig. 5c). Thus, ZnF2 is a DNA-binding domain required for Zld function in type II neuroblasts and dispensable in the early embryo.

## Translational frameshifting generates a truncated Zld proteoform enriched in neuroblasts

While ZnF2 is selectively required in type II neuroblasts, our data did not reveal how this tissue specificity is regulated. One intriguing possibility was revealed by our identification of a unique tissue-enriched Zld polypeptide. The Zld antibody used by us and others was generated against the C-terminus, including ZnF3-6, and recognizes a single polypeptide in both embryos and larval brains enriched for type II neuroblasts (Fig. 4a)^42^. By contrast, immunoblots against a GFP tag on the N-terminus of the endogenously expressed protein identified two polypeptides: one that corresponded to the full-length protein recognized by the anti-Zld antibody and a distinct smaller polypeptide (Fig. 4a). Prior results from two independent laboratories identified a similar species using anti-GFP antibodies, reinforcing our results^43,44^. Compared to the full-length protein, the smaller polypeptide was enriched in type II neuroblasts, whereas the two polypeptides were present at approximately equal levels in stage 5 embryos (Fig. 4a). We confirmed that both polypeptides correspond to Zld as they were recognized by an antibody raised against the N-terminus of Zld (anti-Zld-N) (Extended data Fig 6a). Anti-HA immunoblots on N-terminally HA-tagged Zld expressed from transgenes in type II neuroblasts also identified the same two polypeptides (Extended data Fig. 3c, 6b). As we observed for the endogenous protein, only the larger polypeptide expressed from the transgene was recognized by the anti-Zld-C antibody (Fig. 4a, Extended data Fig. 6b), confirming that the smaller polypeptide did not contain the C-terminal epitope recognized by the anti-Zld-C antibody that includes ZnF3-6. Expression of a cDNA truncated at amino acid 1115 resulted in a protein of approximately the same molecular weight as the smaller product (Extended data Fig. 6b). Thus, we identified an endogenously expressed Zld polypeptide enriched in type II neuroblasts that likely includes ZnF2 but lacks the CAGGTA-binding domain (ZnF3-6) version, raising the possibility that this novel proteoform influences tissue-specific Zld chromatin occupancy.

**Figure 4:**
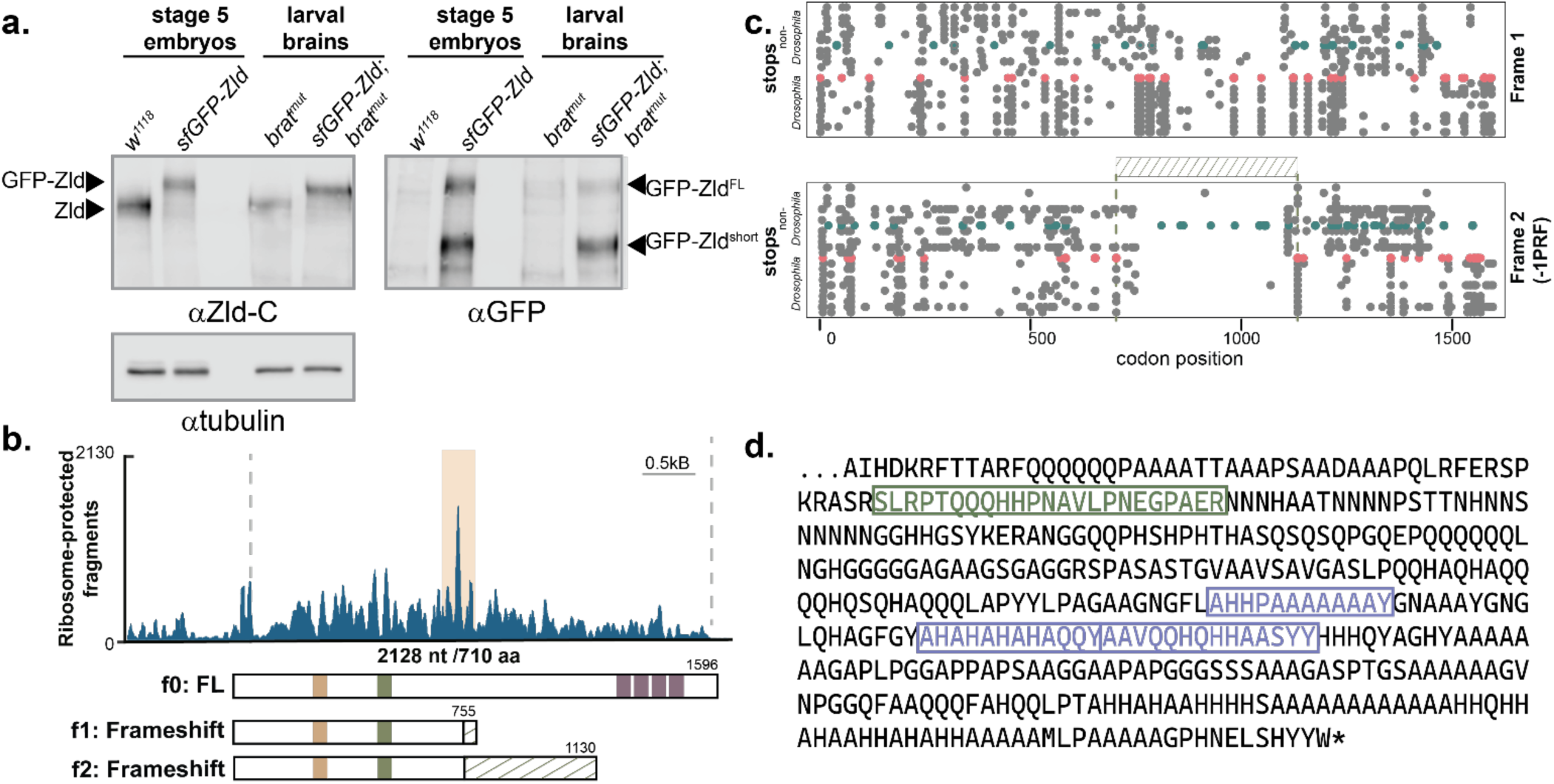
A programmed frame shift generates a truncated Zld proteoform lacking the C-terminal DNA-binding domain. **a.** Immunoblots for GFP, Zld-C (antibody recognizing C terminus) or tubulin on stage 5 embryos or brains of the genotypes indicated. *brat^mut^* is used to enrich for type II neuroblasts. **b.** Ribosome-protected fragments across *zld* from Kronja et al. 2014^49^. Highlighted region (orange) indicates possible ribosome stall site. Schematic models depict the canonical full-length (FL) protein from the original reading frame (f0) and the alternative polypeptides predicted to arise from a frameshifting in both additional frames (f1, f2). Upon frameshifting codon usage is altered producing proteins with distinct C-terminal sequences (green hashed region). **c.** Distribution of stop codons in aligned *zld* coding sequences across alternative reading frames from additional *Drosophila* species as well as other Pancrustacea (see Methods for order of species). Stop codons identified in reading frame 1 (*top*) or reading frame 2 (*bottom*) across all aligned species. Red corresponds to *D. melanogaster* stop codons. Teal indicates *Hyalella azteca*, which encodes a diverged Zld. Hashed region indicates the extended stop-free region in frame 2. **d.** C-terminal amino acids specific to frameshift-derived Zld (frame 2). Peptides identified by mass spectrometry following trypsin (green) or chymotrypsin (purple) digestion are highlighted.

The canonically studied 1596 amino acid Zld protein, Zld-PB, is encoded by a single coding exon. However, an alternatively spliced *zld* transcript, *zld-RD*, which contains a unique downstream exon and an alternative, internal splice junction, is enriched in the central nervous system^45,46^. *zld-RD* encodes a 1373 amino acid protein lacking three of the four C-terminal zinc fingers. This protein is larger than the polypeptide we observed and is recognized by the anti-Zld-C antibody, unlike the shorter proteoform we identified32. Furthermore, we identified the small protein product even when expression was driven from a UAS transgene containing the *zld-RB* open reading frame that lacks the alternative 3’ splice site (Extended data Fig. 3c, 6b). These observations made it highly unlikely that the smaller polypeptide was the result of alternative splicing. Nonetheless, to directly test the role of splicing we generated a silent mutation at the predicted 5’ splice site in the transgene expressing HA-Zld. Immunoblotting confirmed expression of both the full-length and short Zld proteins from this transgene (Extended data Fig. 6b). RNA-seq and single-molecule fluorescent in situ hybridization (FISH) similarly failed to detect *zld-RD* expression in type II neuroblasts^24^. Consistent with these findings, no expression of mCherry-tagged endogenous Zld-PD was detected in type II neuroblasts but was identified in neurons (Extended data Fig. 6c). These data rule out alternative splicing as the mechanism underlying formation of the smaller Zld protein. We next tested the role of proteasome-mediated cleavage. In the *Drosophila* central nervous system, partial proteasomal degradation of the transcription factor Lola results in a truncated product that is essential for sex-specific neuronal wiring and can be detected in S2 cells^47^. Immunoblots with anti-Zld-C antibody on lysate from S2 cells expressing *zld* recognized a single protein product while anti-Zld-N identified two proteins specific to *zld* expression (Extended data Fig. 6d). Similar to the early embryo, both Zld proteins were expressed at approximately equal levels. In contrast to Lola^47^, treatment with the proteasome inhibitor MG132 did not change the ratio between the two Zld proteins (Extended data Fig. 6d), arguing against proteasome-dependent cleavage producing the smaller Zld product.

Having ruled out the most straightforward mechanisms for generating the smaller Zld protein, we hypothesized that it might result from alterations in translation. Analysis of published ribosome profiling in early embryos revealed a pause site in *zld* near the codon for amino acid 710 (aa710), which could result in ribosomal frameshifting (Fig. 4b)^48,49^. Ribosomal frameshifting is best described on viral mRNAs, but it is a highly unusual mechanism on cellular mRNAs^50,51^. While frameshifts normally result in stop codons within the proximal 15-20 amino acids^52^, a −1 shift at this site would create an extended alternative open reading frame that continues for >400 codons. The predicted product would be 1130 amino acids, matching the size of the Zld proteoform enriched in type II neuroblasts (Fig. 4b, Extended data Fig. 7a). The resulting polypeptide would contain ZnF2 but not ZnF3-6 (Fig. 4b). Further support for frameshifting as a potential mechanism to generate this altered proteoform came from evolutionary analysis. Stop codons were purged from the −1 reading frame (frame 2) in this region of *zld* throughout the Pancrustacea lineage, but were present elsewhere in the genome (Fig. 4c)^34^. The conservation of this feature supports the unexpected possibility that the smaller Zld protein results from ribosomal frameshifting.

We exploited the fact that the predicted ribosomal frameshift would create a protein with a unique C-terminus not present in other *Drosophila* proteins to acquire experimental evidence for translation of an alternative reading frame on *zld* mRNA. We included peptides translated from this alternative reading to the search database used for mass spectrometry analysis. We identified four distinct peptides encoded by the alternative reading frame by analyzing immunoprecipitated Zld and bulk embryo lysate using trypsin and chymotrypsin (Fig. 4d, Extended data Fig. 7b). Analysis of published RNA-seq data did not identify transcripts with indels that would be indicative of polymerase slippage (Extended data Fig. 7c)^53^, further supporting translational frameshifting as the mechanism generating the altered Zld proteoform. To experimentally demonstrate that frameshifting resulted in the smaller Zld proteoform, we created the Zld^shortV5^ frameshift reporter by encoding a V5 epitope in the −1 frame just before the predicted stop codon. Importantly, the epitope did not introduce stop codons or encode the V5 epitope in the open reading frame for the canonical, full-length protein. We expressed Zld^shortV5^ in S2 cells or in type II neuroblasts of the larval brain. Both the large and small Zld proteoforms were detected, as expected (Fig. 5a, Extended data Fig. 8a,b). Antibodies against the V5 epitope only recognized a single polypeptide corresponding in size to the smaller proteoform (Fig 5a, Extended data Fig. 8a,b). These data demonstrate that ribosomal frameshifting produces an altered Zld proteoform that lacks the canonical DNA-binding domain (ZnF3-6) while retaining the type II neuroblast-specific DNA-binding domain (ZnF2).

**Figure 5:**
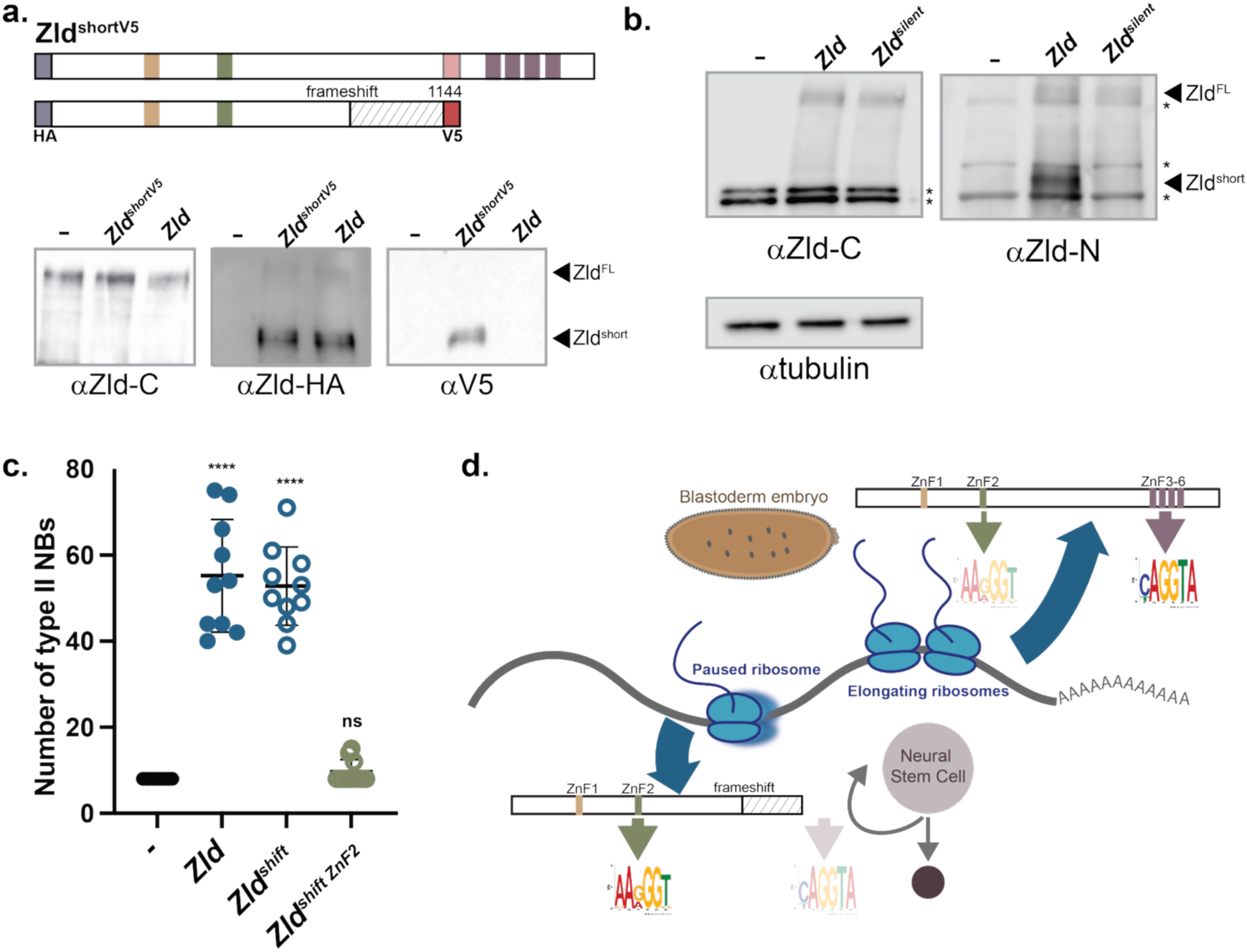
The frameshift-derived Zld proteoform promotes neural stem cell self-renewal in a ZnF2-dependent manner. **a.** Immunoblots for Zld-C, HA, or V5 on *brat^mut^* brains alone or expressing either a wild-type *zld* cDNA or one that encodes a V5-tag in −1 reading frame (Zld^shortV5^). Above, models of predicted proteins produced by Zld^shortV5^, indicating N-terminal HA tag and V5 in frame 2. **b.** Number of type II neuroblasts (NBs) per lobe of brains in which the indicated Zld constructs are driven by *Wor-Gal4, tub-Gal80^ts^*. *Zld^shift^* is a *zld* cDNA that forces expression of the frameshifted product through the addition of an extra nucleotide. Comparisons are to *Wor-Gal4, tub-Gal80^ts^* alone using an ANOVA**. c.** Model of how ribosomal pausing, primarily in type II neuroblasts, leads to an altered proteoform that lacks the CAGGTA-binding C-terminal domain (ZnF3-6) but encodes ZnF2, which promotes binding to AAGGGT-rich sequences enriched in type II neuroblast-specific enhancers. In embryos, frameshifting is less frequent resulting in a protein product that encodes ZnF3-6, which binds CAGGTA-rich embryo-specific enhancers. ****, p < 0.0001. ns = not significant.

To determine whether the region in which we observed pausing of ribosomes was the site of frameshifting, we introduced mutations that were silent in reading frame 0 encoding the full-length protein, but resulted in stop codons in the −1 reading frame (Extended data Fig. 8a,b). Introduction of a stop codon near the predicted frameshift site abolished production of the smaller Zld proteoform (Zld^700–STOP^), whereas a downstream stop codon generated a truncated short product (Zld^1050–STOP^) (Extended data Fig. 8a,b). Neither of these disrupted the expression of the full-length product (Extended data Fig. 8a,b).

These data are consistent with ribosomal pausing while translating ∼710 aa. RNA sequence (codon optimality, RNA structure, RNA modification) or interactions of the nascent polypeptide with the ribosome can induce ribosomal pausing leading to frameshifting^50^. To test these features, we introduced silent mutations in frame 0 that would change the RNA sequence encoding amino acids 701-770 (Zld^silent^) and tested for expression of both Zld proteoforms (Fig. 5b). Expression of Zld^silent^ only produced the canonical, 1596aa Zld proteoform, suggesting that this 210 nt RNA sequence instructs the ribosomal frameshifting that results in the smaller Zld proteoform.

## The frameshift-derived short Zld proteoform is sufficient to promote neuroblast fate

Our identification of the Zld proteoform lacking the canonical DNA-binding domain (ZnF3-6) raised the possibility that this protein occupies previously undetected loci in the embryo. Spike-in normalized ChIP-seq using the GFP epitope on GFP-Zld embryos largely overlapped ChIP-seq using the anti-Zld-C antibody and were enriched for the CAGGTA motif (Extended data Fig. 8c). Peaks unique to either condition were the lowest peaks, suggesting the differences were technical rather than biological. Thus, the smaller Zld proteoform does not strongly occupy unique binding sites in the embryo.

Because ZnF2 is sufficient to promote Zld binding in type II neuroblasts, we tested whether the smaller Zld proteoform is functionally sufficient to promote type II neuroblast-specific activity. To selectively produce the short proteoform, we generated constructs containing an additional nucleotide downstream of the predicted frameshift site to force translation into the alternative reading frame (Zld^shift^) (Extended data Fig. 8d). Zld^shift^ selectively activated the type II neuroblast-specific *tll* reporter, whereas Zld activated both embryonic (*sc*) and type II neuroblast-specific (*tll*) reporters (Extended data Fig. 8e). Mutation of ZnF2 abolished activation by Zld^shift^ (Extended data Fig. 8e). Thus, the 1130aa Zld proteoform is sufficient to activate type II neuroblast, but not embryonic gene expression. To establish the biological role of the smaller proteoform, we tested whether it was sufficient to drive type II neuroblast expansion *in vivo*. Expression of HA-Zld^shift^ in larval brains induced extra type II neuroblasts similarly to full-length Zld, whereas mutation of ZnF2 abolished this activity (Fig. 5b). Our data show that developmentally regulated ribosomal frameshifting regulates tissue-specific binding and activity of the pioneer factor Zld.

## Discussion

The ability of pioneer factors to access their binding motifs in the context of closed chromatin enables them to act at the top of gene-regulatory networks to drive widespread changes in cell identity^13–16^. Nonetheless, these factors show tissue-specific chromatin occupancy. Chromatin structure, co-factor expression and local protein concentration all shape tissue-specific pioneer factor binding^4,5,17,26^. We revealed a new strategy where developmentally programmed ribosomal frameshifting confer tissue-specific binding to the paradigmatic pioneer factor Zld. −1 ribosomal frameshifting produces a short Zld proteoform that is enriched in type II neuroblasts. While this proteoform lacks the canonical DNA-binding domain found in the C-terminus of full-length Zld, we identified a second, previously uncharacterized DNA-binding domain present in both the full-length and frameshifted proteoforms. Frameshifting enables the preferential usage of the two, different DNA-binding domains to govern the distinct Zld-binding profiles in the early embryo and type II neuroblasts.

The deeply conserved second zinc finger of Zld has opposing functions in the embryo and type II neuroblasts. ZnF2 is required for promoting type II neuroblast fate but inhibits Zld-mediated transcriptional activation in the embryo^33^. Our demonstration that ZnF2 encodes a type II neuroblast-specific DNA-binding domain explains these results. ZnF2 directs Zld binding to the *cis*-regulatory elements that drive type II neuroblast fate. These regulatory elements lack the canonical CAGGTA motif bound by Zld in embryos and are instead enriched for the G-rich motif bound by ZnF2. By contrast, the *cis*-regulatory elements that promote zygotic genome activation are highly enriched for the CAGGTA motif bound by ZnF3-6^7,38^. We propose that in the embryo ZnF2 inhibits Zld activity by competing with ZnF3-6 DNA-binding activity, explaining why mutations in ZnF2 lead to hyperactivation of CAGGTA-containing, embryonic genes^33^. This suggests a model in which pioneer activity is regulated by competition between multiple DNA-binding domains, with domain choice dictated by developmental context. This mechanism is unlikely to be unique to Zld. Indeed POU transcription factors, like Oct4, contain multiple DNA-binding domains: both a POU-specific domain and a POU-homeodomain connected by a flexible linker^54,55^. The discovery of a previously unrecognized DNA-binding domain in Zld, despite over a decade of study, underscores the importance of examining pioneer-factor function across developmental contexts. Future insights into the mechanisms governing the tissue-specific usage of distinct binding domains will also impact our understanding of how chimeric oncoproteins that juxtapose multiple DNA-binding to create new transcription factors with emergent and sometimes antagonistic regulatory properties. Together, our data suggest that the modularity provided by multiple DNA-binding domains may allow a flexible mechanism to achieve tissue-specific outcomes and indicate broader principles governing pioneer-factor biology.

We demonstrate that DNA-binding domain use is controlled by developmentally programmed ribosomal frameshifting that results in a Zld proteoform lacking the C-terminal DNA-binding domain. Frame shifting is commonly used in viruses to diversify the proteome, but demonstrated frameshifting on cellular mRNAs is exceedingly rare with only a few other metazoan examples^50,56,57^. Moreover, these few examples appear to be genes with a viral origin, suggesting they retain frameshifting used by the virus and are distinct from the process we report here. Proteoforms generated through frameshifting would not be identified by RNA-sequencing, as they are encoded by the same mRNA, nor by mass spectrometry, that relies on a reference genome. Recent comprehensive analysis of numerous mass spectrometry datasets identified thousands of unique alternative translation events that resulted in single amino acid substitutions, further supporting the potential for the widespread existence of alternative proteoforms^58^. We suggest that stem cells, such as the neuroblasts, may be poised for alternative translational programs as they have low global translation rates despite high levels of ribosome biogenesis^59^.

Numerous mechanisms potentiate ribosomal pausing and frameshifting, including mRNA structure and modifications, tRNA abundance, tRNA modifications, expression of proteins that resolve stalled ribosomes, No-go decay, and levels of ribosome biogenesis^60–67^. Many of these are developmentally regulated. Determining how the ratio of the two Zld proteoforms is controlled will be essential to fully understanding tissue-specific regulation of this pioneer factor, and manipulation of these pathways will enable identification of additional proteins that may be generated through frameshifting.

## Materials and Methods

### Drosophila strains

For CUT&RUN experiments, HA-tagged Zld constructs were expressed in larval brains using the *worniu-Gal4* driver in a *brat* mutant background. Flies carrying *brat¹¹, tub-Gal80ts/CyO, act-GFP* and the indicated *UAS-HA-Zld* transgene were crossed to flies carrying *bratDf, tub-Gal80ts/CyO, act-GFP; worniu-Gal4*. The resulting *brat¹¹/brat^Df^, tub-Gal80ts; UAS-HA-Zld/worniu-Gal4* larvae were raised at 18°C until the third instar (L3) stage and subsequently shifted to 33°C for 24 hours to induce expression of the HA-tagged Zld constructs.

Flies were raised on standard fly food at 25°C (unless otherwise noted). HA-tagged constructs were cloned into pUAST-attB (Addgene). Transgenes were inserted into the *M{3xP3-RFP.attP}ZH-86Fb* docking site using ΦC31 integrase-mediated transgenesis (BestGene, Chino Hills, CA). For ectopic expression of Zld in larval brains, HA-tagged versions of Zld with mutations in ZnF2 (Zld^ZnF2^; C554S), ZnF3-6 (Zld^ZnF3–6^;C1328S, C1371S, C1386S and C1415S); or ZnF2-6 (Zld^ZnF2–6^; all five mutations), as well as full-length Zld containing a V5 tag in the −1 reading frame (Zld^ShortV5^), a single nucleotide insertion in the codon corresponding to amino acid 710 in full-length wild-type (Zld^shift^), or ZnF2-mutant Zld (Zld^shift-ZnF2^) were introduced into *brat*^11^*^/DF^,tub-Gal80^ts^*;*UAS-HA-transgene/Worniu-Gal4* larvae. Larvae were grown at 18°C until they reached L3 stage and then shifted to 33°C for 24 hours to induce ectopic expression of the HA-Zld constructs. GFP-negative and non-Tubby larvae were selected for brains dissections.

### Cell culture, generation of stable cell lines, luciferase assays and RNAi treatment

The stable cell line expressing Zld WT used in this study was previously described^26^. *zld* cDNA containing mutations in ZnF2, ZnF3-6 or ZnF2-6 was cloned into pMT-puro (Addgene) by Gibson assembly (New England Biolabs) as previously described^26^ and cells were continuously selected with 1 µg/ml of puromycin. Zld expression was induced by adding 1000uM CuSO_4_ to the cell culture medium. For luciferases assays, cDNAs were expressed from the *actin* promoter (pAc5.1). cDNAs with mutations in ZnF2 or ZnF3-6 under the *actin* promoter (pAc5.1), as well as full-length Zld containing a V5 tag in the −1 reading frame (Zld^ShortV5^), silent mutations in the 0 reading frame between amino acids 700 and 770 (Zld^silent^), a stop codon in the −1 reading frame near amino acid 700 (Zld^700–STOP^), a stop codon in the −1 reading frame near amino acid 1050 (Zld^1050–STOP^), or a single nucleotide insertion at the codon corresponding to amino acid 710 in full-length wild-type (Zld^shift^) or ZnF2-mutant Zld (Zld^shift-ZnF2^), were used. Cells were plated in 24-well dishes and transiently transfected in triplicate with a total of 300 ng of DNA per well using Effectene transfection reagent (Qiagen), as previously described^32,33^. Luciferase assays were performed using the Dual-Luciferase assay system (Promega). Firefly luciferase activity was normalized to Renilla luciferase activity, and fold activation was calculated relative to the empty-vector control (pAc5.1).

### Immunoblotting

Proteins were transferred to 0.45 μm Immobilon-P PVDF membrane (Millipore, IPVH00010)) in transfer buffer (25 mM Tris, 200 mM Glycine, 20% methanol) for 75 min at 500mA at 4°C. Membranes were blocked with BLOTTO (2.5% non-fat dry milk, 0.5% BSA, 0.5% NP-40, in TBST) for 30 min at room temperature and then incubated with anti-Zld C (1:750; raised against amino acids 1117–1487 of Zld)^42^, anti-Zld N (1:1000; raised against amino acids 100–619 of Zld; this paper), anti-GFP (1:1000) (50430-2-AP, Proteintech), anti-HA–peroxidase (1:500) (clone 3F10, Roche), anti-tubulin (1:5,000) (DM1A, Sigma), anti-V5 (1:1000) (14440-1-AP, Proteintech) overnight at 4 °C. The secondary incubation was performed with goat anti-rabbit IgG-HRP conjugate (1:3000) (Bio-Rad) or anti-mouse IgG-HRP conjugate (1:3000) (Bio-Rad) for 1 hr at room temperature. Blots were treated with SuperSignal West Pico PLUS chemiluminescent substrate (Thermo Fisher Scientific).

### Protein purification

Zld N (Zld^1–619^), Zld N^ZnF2^ (Zld^1–619(ZnF2)^) and Zld C (Zld^1117–1487^) were cloned into pGEX-6P-1, a T7-driven GST expression vector (Addgene). GST-Zld N, GST-Zld N^ZnF2^ and GST-Zld C were purified from *E. coli* in column buffer (20mM Hepes (pH 7.5), 0.5M KCl, 1mM EDTA, 10mM DTT and 0.8mM PMSF and 0.01% NP40) using GST beads (New England Biolabs). After washing, protein was eluted with elution buffer (20mM Hepes (pH 7.6), 0.150 M KCl, 1 mM EDTA, 10 mM DTT and 0.8 mM PMSF, 0.2% NP40, 100% glycerol, and 20mM L-glutathione).

### Electromobility shift assays

EMSAs were performed as previously described^37^. Cy5-labelled probes were incubated with recombinant proteins (Zld N, Zld N^ZnF2^ or Zld C) in buffer containing: 25ng poly[d-(IC)], 12.5mM HEPES, 0.5mM EDTA, 0.5mM EGTA, 5% glycerol, 93.75μM ZnCl2, 0.375mM DTT, 75μM PMSF, 0.075mg/ml BSA, 5mM MgCl2, 0.00625% NP-40 and 50mM KCl at room temperature for 60 min. For EMSAs comparing Zld N and Zld C, 12.5 or 25 nM recombinant protein was incubated with each of three different probes (CAGGTAG, AAGGGTT, GAACTAC). For EMSAs comparing Zld N and Zld N^ZnF2^, 3, 9, or 12.5 nM recombinant protein was incubated with the AAGGGT probe.

The sequence of the oligonucleotides probes that were Cy5 labeled were as follows: GAGAGAGACTACCTGTGGCTCACT (wild type), GAGAGAGAGTAGTTCTGGCTCACT (mutant) and GAGAGAGATTGGGAATGGCTCACT (AAGGGT).

### Protein-binding microarrays

For PBMs, GST-tagged constructs expressing either Zld N^103–608^ and Zld C^1241–1596^ proteins were used as previously described^36^. Briefly, proteins were expressed using the PURExpress In Vitro Protein Synthesis Kit (New England BioLabs) and assayed in duplicate on two PBM arrays containing distinct probe sequences. Brujin sequences were used to construct universal arrays to provide an unbiased representation of all possible 7-mers DNA sequences. Fluorescence intensity data were processed to extract binding signals and used to generate DNA-binding motifs with Top10AlignZ^35^. For quantitative analysis, E-scores and Z-scores were calculated for every 7-mer as previously^35,68^.

### GFP-Zld purification from embryos

*Drosophila* embryos expressing endogenously GFP-tagged Zld^33^ were collected over a 1-hour window and aged for 2 hours to enrich for stage 5 embryos. Embryos were dechorionated in 50% bleach for 90s, washed with distilled water, and homogenized in ice-cold RIPA lysis buffer (10mM Tris-HCl, 0.5mM EDTA, 150mM NaCl, 1% NP-40, 0.5% sodium deoxyclolate, % SDS) supplemented with EDTA-free protease inhibitors (Roche) and 0.8mM PMSF. Lysates were incubated on ice for 30 min with intermittent mixing and clarified by centrifugation at 17,000 × g for 15 min at 4°C. Protein concentration was determined by Bradford assay.

For immunoprecipitation, GFP-Zld was affinity-purified using GFP-Trap magnetic particles (GFP Nano-Trap M-270, ChromoTek) according to the manufacturer’s instructions. Briefly, 25 μL of bead slurry was equilibrated by washing twice with ice-cold bead wash buffer (10mM Tris-HCl, 0.5mM EDTA, 150mM NaCl, 0.05 % NP-40) and then incubated overnight at 4°C with 2 mg of lysate using end-over-end rotation. Beads were collected with a magnetic rack, washed once with wash buffer (10mM Tris-HCl, 0.5mM EDTA, 150mM NaCl), and bound proteins were eluted by boiling the beads for 10 min in 2X SDS loading buffer. Samples were separated by SDS-PAGE, and bands corresponding to full-length and smaller Zld proteoforms were excised from the gel for mass spectrometry analysis.

### Sample preparation for mass spectrometry

For experiments using purified GFP-Zld (n=2), gel slices were destained in a 1:1 mixture of 50 mM ammonium bicarbonate and 100% ethanol for 15 min at 60 °C, with the procedure repeated until the gel pieces became translucent. The pieces were then dehydrated in 100% ethanol for 10 min at room temperature. Proteins were reduced with 20 mM DTT in 50 mM ammonium bicarbonate for 20 min at room temperature and subsequently alkylated with 80 mM chloroacetamide in 50 mM ammonium bicarbonate for 20 min in the dark. Gel pieces were then washed sequentially for 10 min each with 50 mM ammonium bicarbonate, a 1:1 mixture of 50 mM ammonium bicarbonate and 100% ethanol and followed by drying in a vacuum centrifuge. Dried gel pieces were rehydrated with 50 µL 50mM ammonium bicarbonate containing sequencing-grade modified trypsin (Promega) at an approximate enzyme-to-protein ratio of 1:50 and digested overnight at 37 °C. Peptides were recovered by two successive extractions with 100 µL of 80% acetonitrile containing 0.2% formic acid for 10 min at room temperature. The resulting supernatants were pooled, dried to completion in a vacuum concentrator, and reconstituted in 0.2% formic acid prior to LC-MS analysis.

For experiments using stage 5 *w*^1118^ embryos (n=3), embryos were resuspended in denaturing buffer containing 5.4 M guanidine hydrochloride and 100 mM Tris (pH 8.0) and briefly vortexed. Samples were heated at 95 °C for 5 min and probe-sonicated for approximately 10 s. Lysates were centrifuged at 5,000 g for 5 min at room temperature, after which the clarified supernatants were transferred to fresh tubes. Proteins in the clarified supernatants were precipitated by adding methanol to a final concentration of 90% (v/v). The precipitated proteins were collected by centrifugation at 9,000 g for 5 min at RT. The resulting pellets were air-dried in a fume hood for 5 min and resuspended in 8 M urea and 100 mM Tris (pH 8.0).

Aliquots containing 50 µg of protein were supplemented with tris(2-carboxyethyl)phosphine (TCEP) and chloroacetamide to final concentrations of 5 and 20 mM, respectively. Samples were incubated at room temperature for 30 min in the dark. Subsequently, the urea concentration was reduced to 6 M by adding 100 mM Tris (pH 8.0). Chymotrypsin (Promega) was added at a protein-to-enzyme ratio of 30:1 (w/w), and digestion was performed for 3 h at 25 °C with gentle agitation. Digestion was quenched by adding trifluoroacetic acid (TFA) to a final concentration of 0.5% (v/v).

The acidified digests were subjected to solid-phase extraction (SPE), using Sep-Pak C18 SPE cartridges (1 cc, 50 mg; Waters) connected to a vacuum manifold. The cartridges were conditioned with 100% acetonitrile (ACN) and equilibrated by two successive washes with 0.2% formic acid (FA). After loading the acidified digests, the cartridges were washed twice with 0.2% FA, and the retained peptides were eluted with 80% ACN containing 0.2% FA. The eluates were evaporated to dryness in a vacuum concentrator and reconstituted in 0.2% FA before LC-MS analysis.

### LC-MS analysis

Chromatographic separation was performed using a Vanquish Neo UHPLC system directly interfaced with an Orbitrap Astral mass spectrometer via a Nanospray Flex ion source (all from Thermo Fisher Scientific). Peptides were loaded onto an internally packed 40-cm fused-silica capillary column (C18, 1.7 µm particle size, 130 Å pore size), with the column temperature regulated at 50°C by a custom-built heating apparatus. For each analysis, 250 ng of total peptide from *w*^1118^ embryos or 500 ng from GFP-Zld purifications was injected at a constant flow rate of 300 nL/min. The liquid chromatography gradient transitioned from 5% to 46% mobile phase B over a 30-minute active window. Mobile phase A consisted of 0.2% formic acid in water, while mobile phase B comprised 80% acetonitrile containing 0.2% formic acid. Source parameters included a 2 kV spray voltage and an ion-transfer-tube temperature of 280 °C. Mass spectrometry data were collected in data-independent acquisition (DIA) mode. Initial MS1 precursor spectra were acquired utilizing the Orbitrap analyzer at a resolving power of 240,000 across a mass-to-charge (m/z) range of 380-980. The RF lens amplitude was fixed at 40%, with the maximum injection time capped at 50 ms and a normalized automatic gain control (AGC) target of 250%.

For MS2 acquisition, the Astral detector recorded DIA scans across the 380-980 m/z precursor range employing sequential 4 m/z isolation windows without overlap. The instrument was configured with an automated DIA window type and time-based loop control, establishing a rapid, fixed cycle time of 0.6 s to ensure comprehensive precursor coverage. Peptides were fragmented via higher-energy collisional dissociation (HCD) utilizing a normalized collision energy (NCE) of 25%. Product ions were scanned from 150 to 2000 m/z with an RF lens setting of 40%, a maximum injection time of 3.5 ms, and a normalized AGC target of 500%.

### MS data analysis

Raw LC–MS data were processed in Spectronaut v19.7 using the directDIA workflow with default settings. Database searching was performed against the *Drosophila melanogaster* reference proteome obtained from UniProt (proteome ID UP000000803), supplemented with the predicted amino acid sequence of the smaller Zld proteoform.

The false discovery rate was controlled at 1% at the peptide-spectrum match, peptide, and protein-group levels. A q-value threshold of 0.01 was applied to precursor identifications. Protein q-value thresholds were set to 0.01 across the experiment and 0.05 within individual runs. Protein inference was performed using the IDPicker algorithm. Relative peptide quantities were derived from MS2 fragment-ion chromatographic peak areas, and protein abundances were calculated using Spectronaut’s automatic label-free quantification procedure. Cross-run normalization, background-noise removal, and interference correction were enabled. No missing-value imputation was performed in Spectronaut. Parameters not specified above were retained at the BGS Factory Settings.

### Identification of stop codons across species

Codon-aligned coding sequences from 24 insect species were analyzed in R using the seqinr package. The analysis included 13 non-*Drosophila* species (*A. gambiae, A. pisum, A. coluzzii, A. merus, B. mori, D. ponderosae, F. arisanus, H. azteca, M. demolitor, N. vitripennis, N. vespilloides, P. machaon,* and *T. castaneum*) and 11 *Drosophila* species (*D. melanogaster, D. virilis, D. simulans, D. ananassae, D. mauritiana, D. mojavensis, D. pseudoobscura, D. santomea, D. sechellia, D. willistoni,* and *D. yakuba*). For each species, gaps were removed from the aligned CDS, and stop codons (TAA, TAG, and TGA) were identified in the two alternative reading frames relative to the annotated coding frame. Stop-codon positions were then mapped to the alignment and expressed as codon coordinates relative to the *D. melanogaster* sequence. The positions of stop codons were visualized across species for each alternative reading frame using base R graphics.

### Cleavage Under Targets & Release Using Nuclease (CUT&RUN)

For CUT&RUN experiments, HA-tagged Zld constructs were expressed in larval brains using the *worniu-Gal4* driver in a *brat* mutant background. Flies carrying *brat¹¹, tub-Gal80ts/CyO, act-GFP* and the indicated *UAS-HA-Zld* transgene were crossed to flies carrying *brat^Df^, tub-Gal80ts/CyO, act-GFP; worniu-Gal4*. The resulting *brat¹¹/brat^Df^, tub-Gal80ts; UAS-HA-Zld/worniu-Gal4* larvae were raised at 18°C and subsequently shifted to 33°C for 24 hours to induce expression of the HA-tagged Zld constructs.

Brains were dissected from fifty GFP-, non-Tubby (*brat¹¹/brat^Df^, tub-Gal80ts; UAS-HA-Zld/worniu-Gal4)* larvae into Schneider’s medium, transferred into Wash Buffer (20 mM HEPES, 150mM NaCl and 0.5mM spermidine) supplemented with 0.1% BSA as described in CUTANA CUT&RUN (Epicypher) and dounced ten times using a tight pestle. The Epicypher protocol was followed with the following modifications: the Digitonin Buffer was prepared at 0.05% concentration; DNA cleanup was performed using the MinElute Reaction Cleanup Kit (Qiagen); and libraries were prepared using the NEBNext Ultra II DNA Library Prep Kit (E7645S, New England BioLabs), with adaptors diluted 1:30. AMPure XP beads (Beckman Coulter) were used at 1.1X ratio following adaptor ligation and 0.9X following PCR. PCR cycling parameters followed those outlined in the Epicypher protocol. Antibodies used included 1µL of anti-Zld^42^ and 1µL of anti-HA (12CA5; Sigma-Aldrich). Libraries were sequenced on an Illumina NovaSeq 6000 platform using 150 bp paired-end reads at the UW-Madison Biotechnology Center.

### CUT&RUN Analysis

Adapters and low-quality bases were removed using Trimmomatic-0.39^69^. Reads were aligned to the *Drosophila melanogaster* reference genome (version dm6) using bowtie 2 v2.3.5^70^. Unmapped, multiply aligning, mitochondrial, and scaffold reads were removed. To identify regions that were enriched in immunoprecipitated samples relative to input controls, peak calling was performed using MACS v2^71^ with the following parameters: −g 1.2e8, −-call-summits. From experiments using Zld antibody peaks were called with respect to a Rab-IgG CUT&RUN control. From experiments using HA antibody peaks were called with respect to a HA CUT&RUN experiment in a *brat* mutant brain (no transgene overexpression). To focus analysis on robust, high-quality peaks, we used 100 bp up- and downstream of peak summits, and regions belonging to contigs and unmapped chromosomes were removed. z score-normalized bigWigs were created by subtracting the mean read coverage (counts) from merged replicate read counts in 10bp bins across the entire genome and dividing by the standard deviation. Heatmaps and average signal line plots were generated using deepTools2 ^72^ with z score-normalized bigWig files.

### ChIP-seq

ChIP-seq experiments were performed on hand-sorted stage 5 embryos laid by *zld*^ZnF2^ homozygous females, as previously described^33^, *w*^1118^ embryos as a wild-type control, and GFP-Zld embryos. For each genotype, two biological replicates were collected, with one thousand embryos per replicate. Briefly, embryos were dechorionated in 50% bleach for 3 minutes and fixed in 0.45% formaldehyde and homogenized in RIPA buffer (50 mM Tris-Cl pH 8.0, 0.1% SDS, 1% Triton-100, 0.5% deoxycholate and 150 mM NaCl). All subsequent steps were performed as previously described^53,73^. For *w*^1118^ and *zld*^ZnF2^, reactions included a spike-in (5%) of *Drosophila virilis* fixed and sonicated chromatin. For normalization of GFP-Zld ChIP-seq, chromatin prepared from mouse cells expressing an H3.3-GFP fusion protein was added to *Drosophila* chromatin at a 1:750 ratio. 2% of the supernatant was set aside as input, and the remainder was incubated at 4 °C overnight with antibodies (7 µl anti-Zld and 6 µg anti-GFP (Abcam 290)). For each ChIP-seq experiment in S2 cells, ∼25 million cells were cross-linked and processed following the protocol previously described^26^. 2% of the supernatant was set aside as input, and the remainder was incubated at 4 °C overnight with antibodies (7 µl anti-Zld). For each genotype, two biological replicates were collected. Preparation of sequencing libraries of both ChIP-seq experiments in embryos and S2 cells was performed using the NEBNext Ultra II kit (NEB) with eight PCR cycles for library amplification. Sequencing was performed on an Illumina Novaseq 6000 instrument using 50-bp single-end reads.

### ChIP-seq analysis

Adapters and low-quality bases were removed using Trimmomatic-0.39^69^. Reads were aligned to the *Drosophila melanogaster* reference genome (version dm6) using bowtie 2 v2.3.5^70^. Unmapped, multiply aligning, mitochondrial, and scaffold reads were removed. To identify regions that were enriched in immunoprecipitated samples relative to input controls, peak calling was performed using MACS v2^71^ with the following parameters: −g 1.2e8, −-call-summits. The BEDtools intersectBed function was used to define high-confidence peaks present in both replicates^74^. To focus analysis on robust, high-quality peaks, we used 100 bp up- and downstream of peak summits, and regions belonging to contigs and unmapped chromosomes were removed. All bigWigs were generated using bamCoverage from deepTools v3.5.192. For ChIP-seq experiment in S2 cells z-score normalized bigWigs were created by taking the average read count across all bins genome wide and dividing by the standard deviation. IPs for Zld in the embryo were spike-in normalized by a scaling factor that was determined by the ratio of percent *D. virilis* chromatin aligned in the IPs as compared to input. GFP-Zld IPs were spike-in normalized by a scaling factor that was determined by the ratio of percent mouse H3.3-GFP chromatin aligned in the IPs as compared to input. Heatmaps and average signal line plots were generated using deepTools2^72^ with z score-normalized (for S2 cells) or spike-in normalized (for embryos) bigWig files.

### ATAC-seq

For each ATAC-seq experiment in S2 cells, ∼200 thousand cells were processed following the protocol previously described^26^. For each genotype, two biological replicates were collected. Samples were tagmented with 2.5 µl of Tagment DNA Enzyme (Illumina) and incubated at 37 °C for 30 min. After DNA tagmentation, DNA was purified using the MinElute Reaction Cleanup kit (Qiagen) and eluted in 10 µl of buffer EB. DNA was PCR amplified for twelve cycles. Amplified libraries were purified using a 1.2× ratio of Axygen paramagnetic beads. Sequencing was performed on an Illumina NovaSeq X Plus instrument using 150-bp pair-end reads.

### ATAC-seq analysis

Adaptor sequences were removed from raw ATAC-seq reads using NGmerge (v0.3)^75^. The resulting reads were aligned to the *D. melanogaster* genome (dm6) with Bowtie2 using −k 2 −-very-sensitive −-no-mixed −-no-discordant −X 5000^70^. Reads with a mapping quality below 30, as well as those mapping to the mitochondrial genome or scaffolds, were excluded from further analysis. As described previously, only fragments <100 bp in length were considered for downstream analysis^26^. Reads from all replicates were merged, and MACS2 was employed for peak calling with −f BAMPE −-keep-dup all −g dm −-call-summits^76^. featureCounts was used to quantify reads within 200 bp of peak summits (±100 bp)^77^. Differential chromatin accessibility was determined using DESeq2, and regions with an adjusted *P* value <0.05 were considered significantly differentially accessible^78^.

### Immunostaining and quantification of type II neuroblasts

Third instar larval brains were dissected in solution containing 4% formaldehyde, 1xPBS and fixed in 100 mM PIPES (pH6.9), 1 mM EGTA, 0.3 % Triton X-100, and 1 mM MgSO4 for 23 min at room temperature. Fixed brains were washed with 0.3 % Triton X-100 in 1x PBS (PBST). After removing fix solution, samples were incubated with primary antibodies for 3 hrs at room temperature. Samples were washed with PBST and incubated with secondary antibodies and Phalloidin (ThermoFisher Scientific) overnight at 4°C. Samples were washed with PBST and equilibrated in Prolong Gold antifade mount (ThermoFisher Scientific). Brains were mounted with dorsal surface up. Confocal images were acquired on a Leica SP5 scanning confocal microscope (Leica Microsystems Inc). To quantify the number of type II neuroblasts, images of brain lobes were sequentially taken at every 1.5 mm from dorsal to ventral. Ten brains per genotype were used to obtain data.

## Data availability

The data that support this study are available from the corresponding authors upon reasonable request. Sequencing data for CUT&RUN from larval brains have been deposited in GEO under accession code GSE308535. ATAC-seq data generated in S2 cells are under accession code GSE308658. ChIP-seq data are under accession codes GSE308659 and GSE349870. bigWig files are also available under each accession code. Previously published sequencing data for ATAC-seq in the larval brain are under accession code GSE150931 and for ribosome footprinting in the early embryo under accession code GSE52799. Mass spectrometry data have been deposited in MassIVE under accession number MSV000103630.

## Acknowledgements

Stocks obtained from the Bloomington *Drosophila* Stock Center (National Institutes of Health P40OD018537) were used in this study. Flybase (FB2024_02) was used throughout the study.

## Author Contributions

ETZ, MMH, HK, CYL, AAM and CAR conceived the study. Most experiments were designed and analyzed by ETZ, HK, and MMH. H-YL, CAR, AY, and TRH performed protein-binding array analysis. MM, BC, and JJC performed and analyzed mass spectrometry. EDL generated and analyzed a subset transgenic fly lines. ETZ and MMH wrote the original draft of the manuscript. All authors reviewed and edited the manuscript. MMH, CYL, AAM, JJC, CAR and TRH acquired funding.

## Reference

1. Buecker, C. et al. Reorganization of Enhancer Patterns in Transition from Naive to Primed Pluripotency. Cell Stem Cell 14, 838–853 (2014).

2. Chronis, C. et al. Cooperative Binding of Transcription Factors Orchestrates Reprogramming. Cell 168, 442–459.e20 (2017).

3. Donaghey, J. et al. Genetic determinants and epigenetic effects of pioneer-factor occupancy. Nat Genet 50, 250–258 (2018).

4. Soufi, A., Donahue, G. & Zaret, K. S. Facilitators and Impediments of the Pluripotency Reprogramming Factors’ Initial Engagement with the Genome. Cell 151, 994–1004 (2012).

5. Mayran, A. et al. Pioneer factor Pax7 deploys a stable enhancer repertoire for specification of cell fate. Nat Genet 50, 259–269 (2018).

6. Larson, E. D. et al. Cell-type-specific chromatin occupancy by the pioneer factor Zelda drives key developmental transitions in Drosophila. Nat Commun 12, 7153 (2021).

7. Harrison, M. M., Li, X.-Y. Y., Kaplan, T., Botchan, M. R. & Eisen, M. B. Zelda binding in the early Drosophila melanogaster embryo marks regions subsequently activated at the maternal-to-zygotic transition. PLoS Genet 7, e1002266 (2011).

8. Liang, H. L. et al. The zinc-finger protein Zelda is a key activator of the early zygotic genome in Drosophila. Nature 456, 400–403 (2008).

9. Schulz, K. N. et al. Zelda is differentially required for chromatin accessibility, transcription factor binding, and gene expression in the early *Drosophila* embryo. Genome Research 25, 1715–1726 (2015).

10. Sun, Y. et al. Zelda overcomes the high intrinsic nucleosome barrier at enhancers during Drosophila zygotic genome activation. Genome research 25, 1703–14 (2015).

11. Iwafuchi-Doi, M. & Zaret, K. S. Pioneer transcription factors in cell reprogramming. Genes Dev. 28, 2679–2692 (2014).

12. Zaret, K. S. Pioneer Transcription Factors Initiating Gene Network Changes. Annu Rev Genet 54, 367–385 (2020).

13. Balsalobre, A. & Drouin, J. Pioneer factors as master regulators of the epigenome and cell fate. Nat Rev Mol Cell Biol 23, 449–464 (2022).

14. Barral, A. & Zaret, K. S. Pioneer factors: roles and their regulation in development. Trends in Genetics 40, 134–148 (2024).

15. Horisawa, K. & Suzuki, A. The role of pioneer transcription factors in the induction of direct cellular reprogramming. Regenerative Therapy 24, 112–116 (2023).

16. Larson, E. D., Marsh, A. J. & Harrison, M. M. Pioneering the developmental frontier. Molecular Cell 81, 1640–1650 (2021).

17. Freund, M. M., Harrison, M. M. & Torres-Zelada, E. F. Exploring the reciprocity between pioneer factors and development. Development 151, dev201921 (2024).

18. Schulz, K. N. & Harrison, M. M. Mechanisms regulating zygotic genome activation. Nature Reviews Genetics 20, 221–234 (2019).

19. Vastenhouw, N. L., Cao, W. X. & Lipshitz, H. D. The maternal-to-zygotic transition revisited. Development 146, dev161471 (2019).

20. Kobayashi, W. & Tachibana, K. Awakening of the zygotic genome by pioneer transcription factors. Current Opinion in Structural Biology 71, 94–100 (2021).

21. Yartseva, V. & Giraldez, A. J. The Maternal-to-Zygotic Transition During Vertebrate Development: A Model for Reprogramming. Curr Top Dev Biol 113, 191–232 (2015).

22. Rajan, A., Ostgaard, C. M. & Lee, C.-Y. Regulation of Neural Stem Cell Competency and Commitment during Indirect Neurogenesis. Int J Mol Sci 22, 12871 (2021).

23. Homem, C. C. F. & Knoblich, J. A. Drosophila neuroblasts: a model for stem cell biology. Development 139, 4297–4310 (2012).

24. Reichardt, I. et al. The tumor suppressor Brat controls neuronal stem cell lineages by inhibiting Deadpan and Zelda. EMBO Reports 19, 102–117 (2018).

25. Andersson, R. & Sandelin, A. Determinants of enhancer and promoter activities of regulatory elements. Nat Rev Genet 21, 71–87 (2020).

26. Gibson, T. J., Larson, E. D. & Harrison, M. M. Protein-intrinsic properties and context-dependent effects regulate pioneer factor binding and function. Nat Struct Mol Biol 31, 548–558 (2024).

27. Heinz, S., Romanoski, C. E., Benner, C. & Glass, C. K. The selection and function of cell type-specific enhancers. Nat Rev Mol Cell Biol 16, 144–154 (2015).

28. Rada-Iglesias, A. et al. A unique chromatin signature uncovers early developmental enhancers in humans. Nature 470, 279–283 (2011).

29. Zhang, J. et al. Distinct H3K9me3 heterochromatin maintenance dynamics govern different gene programmes and repeats in pluripotent cells. Nat Cell Biol 26, 2115–2128 (2024).

30. Ferrie, J. J., Karr, J. P., Tjian, R. & Darzacq, X. “Structure”-function relationships in eukaryotic transcription factors: The role of intrinsically disordered regions in gene regulation. Molecular Cell 82, 3970–3984 (2022).

31. Jonas, F., Navon, Y. & Barkai, N. Intrinsically disordered regions as facilitators of the transcription factor target search. Nat Rev Genet 26, 424–435 (2025).

32. Hamm, D. C., Bondra, E. R. & Harrison, M. M. Transcriptional Activation Is a Conserved Feature of the Early Embryonic Factor Zelda That Requires a Cluster of Four Zinc Fingers for DNA Binding and a Low-complexity Activation Domain. Journal of Biological Chemistry 290, 3508–3518 (2015).

33. Hamm, D. C. et al. A conserved maternal-specific repressive domain in Zelda revealed by Cas9-mediated mutagenesis in Drosophila melanogaster. PLoS Genet 13, e1007120 (2017).

34. Ribeiro, L. et al. Evolution and multiple roles of the Pancrustacea specific transcription factor zelda in insects. PLOS Genetics 13, e1006868 (2017).

35. Weirauch, M. T. et al. Determination and Inference of Eukaryotic Transcription Factor Sequence Specificity. Cell 158, 1431–1443 (2014).

36. Brennan, K. J. et al. Chromatin accessibility in the Drosophila embryo is determined by transcription factor pioneering and enhancer activation. Developmental Cell 58, 1898–1916.e9 (2023).

37. McDaniel, S. L. et al. Continued Activity of the Pioneer Factor Zelda Is Required to Drive Zygotic Genome Activation. Molecular Cell 74, 185–195.e4 (2019).

38. Nien, C.-Y. et al. Temporal Coordination of Gene Networks by Zelda in the Early Drosophila Embryo. PLoS Genet 7, e1002339 (2011).

39. Fallacaro, S. et al. A fine kinetic balance of interactions directs transcription factor hubs to genes. bioRxiv 2024.04.16.589811 (2024) doi:10.1101/2024.04.16.589811.

40. Janssens, D. H. et al. Earmuff restricts progenitor cell potential by attenuating the competence to respond to self-renewal factors. Development 141, 1036–1046 (2014).

41. Sadowski, I., Ma, J., Triezenberg, S. & Ptashne, M. GAL4-VP16 is an unusually potent transcriptional activator. Nature 335, 563–564 (1988).

42. Harrison, M. M., Botchan, M. R. & Cline, T. W. Grainyhead and Zelda compete for binding to the promoters of the earliest-expressed Drosophila genes. Developmental biology 345, 248–255 (2010).

43. Galouzis, C. C. et al. Chip (Ldb1) is a putative cofactor of Zelda forming a functional bridge to CBP during zygotic genome activation. Molecular Cell 85, 2425–2441.e9 (2025).

44. Li, X.-Y. & Eisen, M. B. The Zelda Interactome Reveals Diverse Co-factors Essential for Pioneer Factor-Mediated Zygotic Genome Activation. Preprint at 10.64898/2026.02.23.707495 (2026).

45. Giannios, P. & Tsitilou, S. G. The embryonic transcription factor Zelda of Drosophila melanogaster is also expressed in larvae and may regulate developmentally important genes. Biochemical and Biophysical Research Communications 438, 329–333 (2013).

46. Pearson, J. C., Watson, J. D. & Crews, S. T. Drosophila melanogaster Zelda and Single-minded collaborate to regulate an evolutionarily dynamic CNS midline cell enhancer. Developmental Biology 366, 420–432 (2012).

47. Sato, K., Ito, H., Yokoyama, A., Toba, G. & Yamamoto, D. Partial proteasomal degradation of Lola triggers the male-to-female switch of a dimorphic courtship circuit. Nat Commun 10, 166 (2019).

48. Dinman, J. D. Mechanisms and implications of programmed translational frameshifting. WIREs RNA 3, 661–673 (2012).

49. Kronja, I. et al. Widespread Changes in the Posttranscriptional Landscape at the Drosophila Oocyte-to-Embryo Transition. Cell Reports 7, 1495–1508 (2014).

50. Atkins, J. F., Loughran, G., Bhatt, P. R., Firth, A. E. & Baranov, P. V. Ribosomal frameshifting and transcriptional slippage: From genetic steganography and cryptography to adventitious use. Nucleic Acids Res gkw530 (2016) doi:10.1093/nar/gkw530.

51. Hill, C. H. & Brierley, I. Structural and Functional Insights into Viral Programmed Ribosomal Frameshifting. Annual Review of Virology 10, 217–242 (2023).

52. Mulroney, T. E. et al. N1-methylpseudouridylation of mRNA causes +1 ribosomal frameshifting. Nature 625, 189–194 (2024).

53. Gaskill, M. M., Gibson, T. J., Larson, E. D. & Harrison, M. M. GAF is essential for zygotic genome activation and chromatin accessibility in the early Drosophila embryo. eLife 10, e66668 (2021).

54. Carminati, M., Vecchia, L., Stoos, L. & Thomä, N. H. Pioneer factors: Emerging rules of engagement for transcription factors on chromatinized DNA. Current Opinion in Structural Biology 88, 102875 (2024).

55. Malik, V., Zimmer, D. & Jauch, R. Diversity among POU transcription factors in chromatin recognition and cell fate reprogramming. Cell Mol Life Sci 75, 1587–1612 (2018).

56. Firth, A. E. & Brierley, I. Non-canonical translation in RNA viruses. Journal of General Virology 93, 1385–1409 (2012).

57. Loughran, G. et al. Programmed ribosomal frameshifting during *PLEKHM2* mRNA decoding generates a constitutively active proteoform that supports myocardial function. Sci. Adv. 11, eady1742 (2025).

58. Tsour, S. et al. Alternate RNA decoding results in stable and abundant proteins in mammals. Nature 656, 506–515 (2026).

59. Gao, Y. et al. Emerging roles of ribosome translation in stem cells and stem cell therapy - a review. Cell Biosci 15, 71 (2025).

60. Brierley, I. & Pennell, S. Structure and Function of the Stimulatory RNAs Involved in Programmed Eukaryotic −1 Ribosomal Frameshifting. Cold Spring Harbor Symposia on Quantitative Biology 66, 233–248 (2001).

61. Caliskan, N. et al. Conditional Switch between Frameshifting Regimes upon Translation of dnaX mRNA. Molecular Cell 66, 558–567.e4 (2017).

62. Chang, W. D. & Choe, Y.-J. Quality control and signaling pathways at stalled ribosomes. Exp Mol Med 58, 82–93 (2026).

63. Dulude, D. Characterization of the frameshift stimulatory signal controlling a programmed −1 ribosomal frameshift in the human immunodeficiency virus type 1. Nucleic Acids Research 30, 5094–5102 (2002).

64. Giedroc, D. P. & Cornish, P. V. Frameshifting RNA pseudoknots: Structure and mechanism. Virus Research 139, 193–208 (2009).

65. Li, Y., et al. Transactivation of programmed ribosomal frameshifting by a viral protein. Proc. Natl. Acad. Sci. U.S.A. 111, (2014).

66. Matsufuji, S. et al. Autoregulatory frameshifting in decoding mammalian ornithine decarboxylase antizyme. Cell 80, 51–60 (1995).

67. Urbonavic-Ius, J., et al. Transfer RNA modifications that alter +1 frameshifting in general fail to affect −1 frameshifting. RNA 9, 760–768 (2003).

68. Berger, M. F. et al. Compact, universal DNA microarrays to comprehensively determine transcription-factor binding site specificities. Nat Biotechnol 24, 1429–1435 (2006).

69. Bolger, A. M., Lohse, M. & Usadel, B. Trimmomatic: a flexible trimmer for Illumina sequence data. Bioinformatics 30, 2114–2120 (2014).

70. Lawrence, M. et al. Software for Computing and Annotating Genomic Ranges. PLoS Comput Biol 9, e1003118 (2013).

71. Zhang, Y. et al. Model-based Analysis of ChIP-Seq (MACS). Genome Biol 9, R137 (2008).

72. Ramírez, F. et al. deepTools2: a next generation web server for deep-sequencing data analysis. Nucleic acids research 44, W160–W165 (2016).

73. Marsh, A. J. et al. Catalytic-dependent and independent functions of the histone acetyltransferase CBP promote pioneer-factor-mediated zygotic genome activation. Molecular Cell 85, 2409–2424.e8 (2025).

74. Quinlan, A. R. & Hall, I. M. BEDTools: a flexible suite of utilities for comparing genomic features. Bioinformatics 26, 841–842 (2010).

75. Gaspar, J. M. NGmerge: merging paired-end reads via novel empirically-derived models of sequencing errors. BMC Bioinformatics 19, 536 (2018).

76. Zhang, Y. et al. Model-based Analysis of ChIP-Seq (MACS). Genome Biol 9, R137 (2008).

77. Liao, Y., Smyth, G. K. & Shi, W. featureCounts: an efficient general purpose program for assigning sequence reads to genomic features. Bioinformatics 30, 923–930 (2014).

78. Love, M. I., Huber, W. & Anders, S. Moderated estimation of fold change and dispersion for RNA-seq data with DESeq2. Genome Biol 15, 550 (2014).

