## Supplemental Figures for "Programmed ribosomal frameshifting regulates tissue-specific pioneer activity"

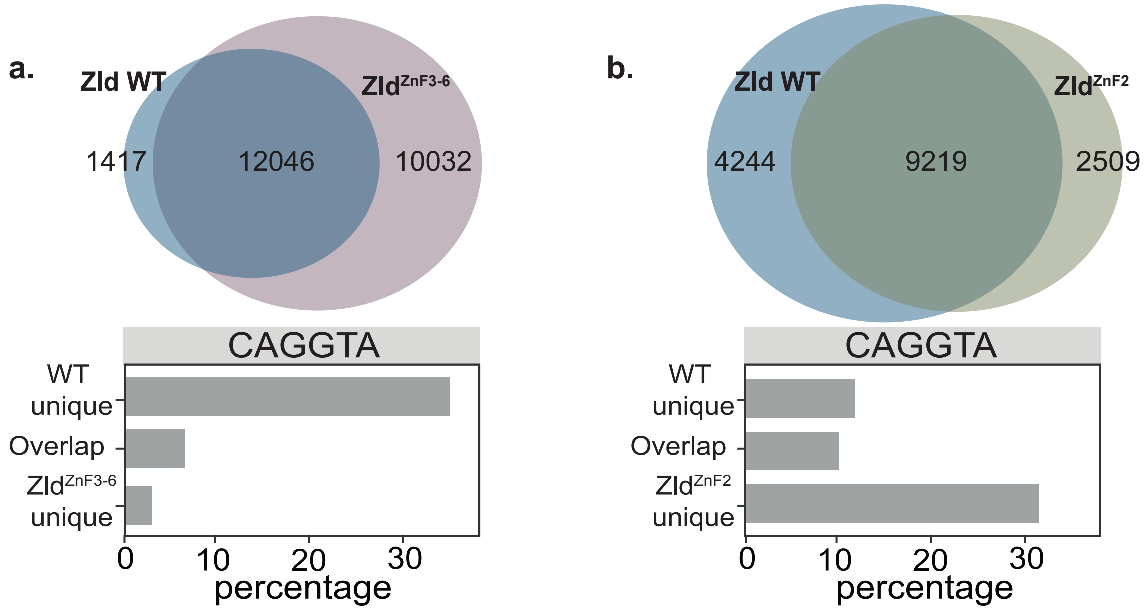


**Extended Data Fig. 1: Mutations in ZnF2 promote Zld binding to CAGGTA-containing sequences**. **a.** Venn diagram of all ChIP-seq peaks for wild-type Zld (Zld WT) and Zld$^{ZnF3-6}$ (*above*). Percentage of peaks that contain the canonical Zld-binding motif, CAGGTA, in each category (*below*). **b.** Venn diagram of all ChIP-seq peaks for Zld WT and Zld$^{ZnF2}$ (*above*). Percentage of peaks that contain the canonical Zld-binding motif, CAGGTA, in each category (*below*).

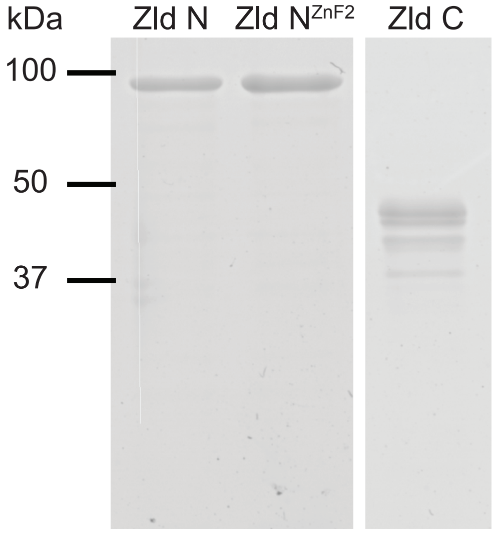


**Extended Data Fig. 2: Purity of recombinant Zld fragments used for domain-specific biochemical analyses.** Coomassie-stained SDS-PAGE gel showing purified recombinant Zld proteins used for EMSAs. Proteins are GST tagged, which was used for purification following expression in *E. coli*.

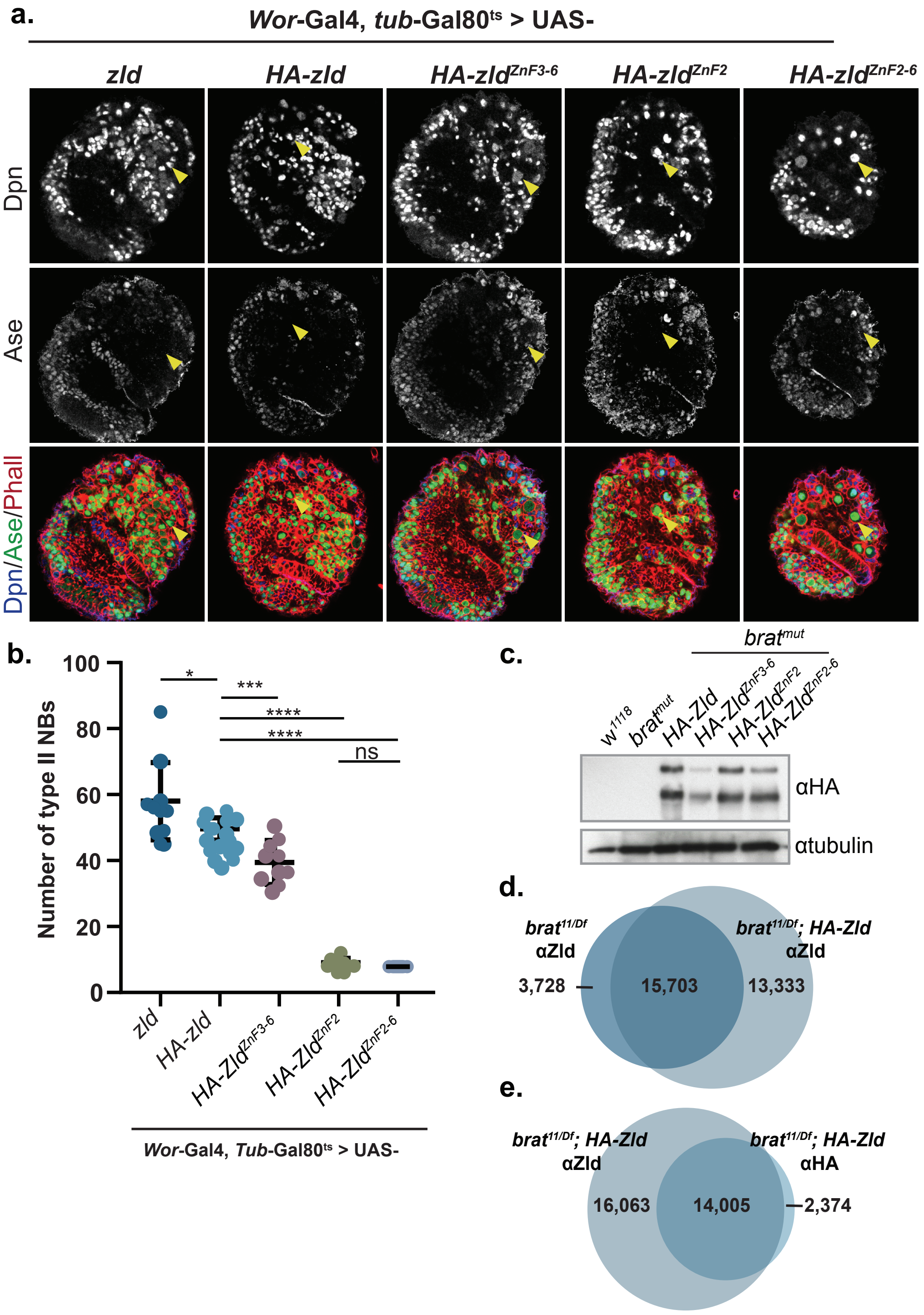


**Extended Data Fig. 3: The N-terminal HA tag does not disrupt Zld binding or activity when expressed in neuroblasts. a.** Immunostaining of third instar larval brains lobes expressing ectopic Zld or HA-Zld constructs as indicated. Expression is driven in type I and type II neuroblasts by *Wor-Gal4, tub-Gal80$^{ts}$*. Yellow arrows indicate a selection of Dpn+, Ase- type II neuroblasts. Scale bar, 20μm. **b.** Number of type II neuroblasts per lobe of brains in which the indicated Zld or HA-Zld constructs are expressed. For each genotype: n = 10 brains. **, $p < 0.01$, , ***, $p < 0.001$, ****, $p < 0.0001$, ns= not significant as determined by ANOVA. **c.** Immunoblots for HA-tagged proteins in brains dissected from the genotypes indicated. Tubulin is shown as a loading control. **d.** Venn diagram of anti-Zld CUT&RUN peaks from *brat$^{11/Df}$* brains and *brat$^{11/Df}$* brains expressing HA-Zld. **e.** Venn diagram of anti-Zld CUT&RUN peaks and anti-HA CUT&RUN peaks from *brat$^{11/Df}$* brains expressing HA-Zld.

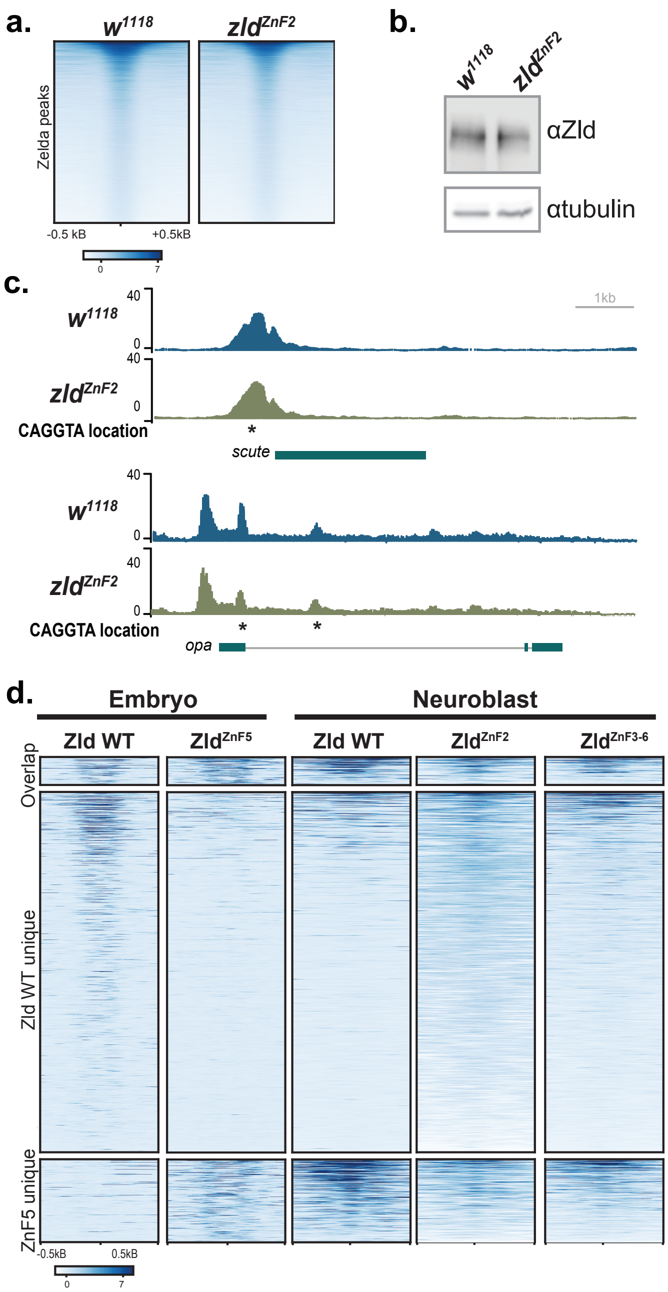


**Extended Data Fig. 4: ZnF2 is dispensable for Zld binding in the embryo, but may promote redistribution when ZnF5 is mutated. a.** Heatmaps of Zld ChIP-seq from embryos expressing $Zld^{WT}$ ($w^{1118}$) or $Zld^{ZnF2}$ centered on Zld-bound peaks. **b.** Immunoblot for Zld on stage 5 embryos demonstrate equivalent protein levels for wild type and ZnF2 mutant. Tubulin is shown as a loading control.**c.** Genome browser tracks showing representative loci of up-regulated Zld-target genes from Hamm et al. *PLoS Genet* 2017. Zld binding in $w^{1118}$ and $zld^{ZnF2}$ embryos along with CAGGTA motifs are shown. **d.** Heatmaps of Zld occupancy in embryos and neuroblasts for the genotypes indicated above at genomic sites defined by comparing wild-type Zld and $Zld^{ZnF5}$ in the embryo (from Fallacaro et al. bioRxiv 2024).

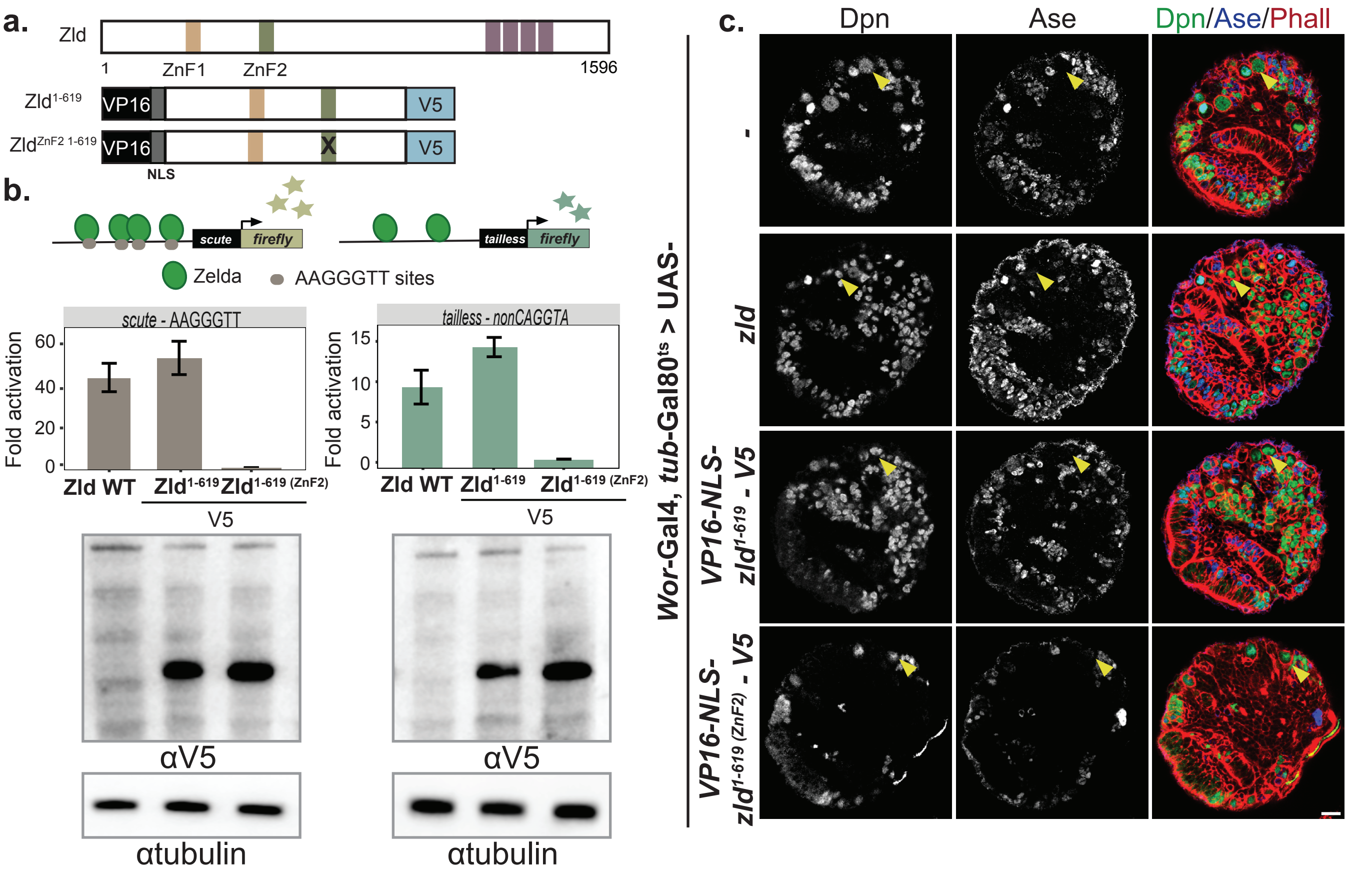


**Extended Data Fig. 5: The N-terminus of Zld is sufficient to promote binding to neuroblast-target genes. a.** Cartoon schematic of VP16-NLS-Zld constructs. **b.** Luciferase reporter assays with full-length Zld, Zld$^{1\text{-}619}$ WT and ZnF2 mutant (Zld$^{1\text{-}619\ (ZnF2)}$). Data are shown for both the *scute* reporter with the four CAGGTA replaced with AAGGGT and the neuroblast-specific *tll* enhancer. Immunoblots for V5 tagged proteins indicate protein expression levels. Tubulin is shown as a loading control. **c.** Immunostaining of third instar larval brains lobes expressing ectopic Zld or VP16-NLS constructs as indicated. Expression is driven in type I and type II neuroblasts by *Wor-Gal4, tub-Gal80$^{ts}$*. Yellow arrows indicate a selection of Dpn+, Ase- type II neuroblasts. Scale bar, 20μm.

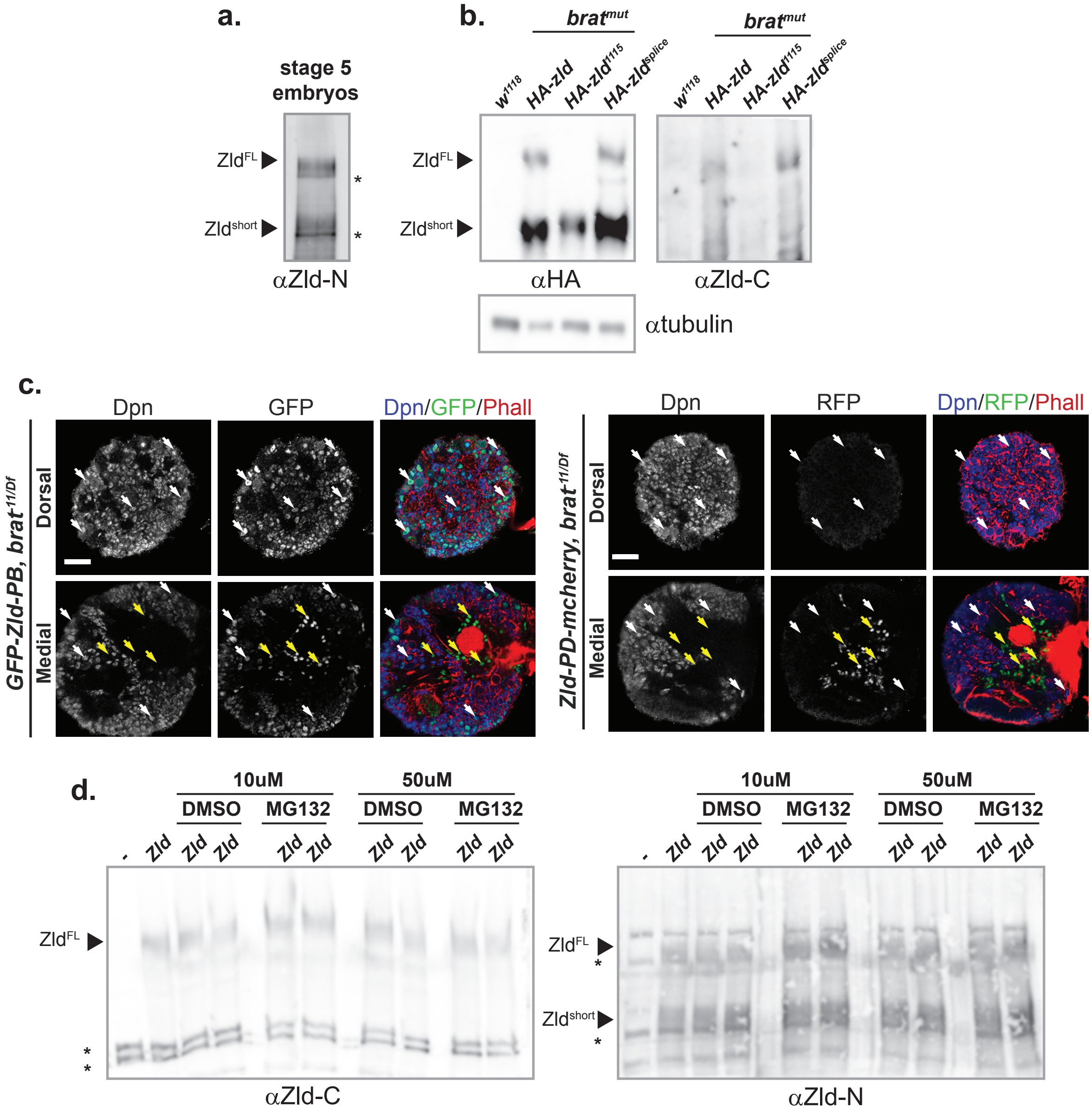


**Extended Data Fig. 6: The truncated Zld proteoform is not generated by splicing or proteolytic cleavage. a.** Immunoblot for Zld on stage 5 embryos with antibodies raised against the N-terminus recognizes two polypetides. The two Zld proteoforms are indicated on the left. *, nonspecific crossreactivity **b.** Immunoblots for HA, Zld (C-term) and tubulin from HA-tagged Zld in brains dissected from the genotypes indicated. Tubulin is shown as a loading control. **c**. Immunostaining of third instar larval brains lobes expressing endogenous GFP-Zld-PB or Zld-PD-mcherry. White arrows indicate Dpn+, neuroblasts and Yellow arrows indicate Dpn-, neurons. Scale bar, 50 µm. **d.** Immunoblot for Zld with antibodies against the C-terminus (Zld-C) or N-terminus (Zld-N) in cells treated with the proteosome inhibitor MG132 (at the concentrations indicated) or DMSO as a control. The two Zld proteoforms are indicated on the left. *, nonspecific crossreactivity a.

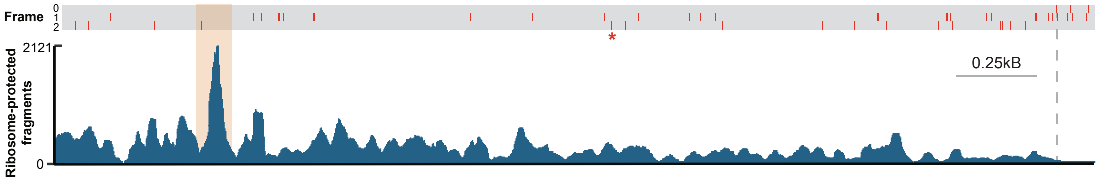


b.

| Frameshift-derived Zld short-specific C-terminal (701-1130 aa) peptides | position (aa) | q-value |
|---|---|---|
| SLRPTQQQHHPNAVLPNEGPAER | 764 - 786 | $1.6 \times 10^{-7}$ |
| AHHPAAAAAAAY | 931 - 941 | $3.8 \times 10^{-6}$ |
| AHAHAHAHAQQY | 959 - 970 | $8.5 \times 10^{-7}$ |
| AAVQQHQHHAASYY | 971 - 984 | $6.8 \times 10^{-7}$ |

c.

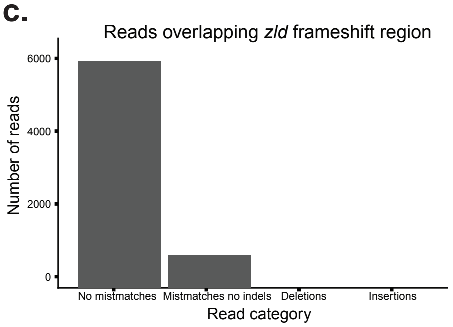


**Extended Data Fig. 7: Ribosome frameshifting generates an alternative Zld proteoform. a.** Ribosome-protected fragment density across the *D. melanogaster zld* coding sequence. Red lines indicate stop codon positions in canonical reading frame (0) and alternative reading frames (frame 1 and 2). * represents stop codon at 1130 aa. **b.** Peptides mapping to the predicted frameshift-derived C-terminal region (aa 701-1130) were identified by DIA mass spectrometry following trypsin (green) or chymotrypsin (purple) digestion. Peptides are supported by high-confidence identifications (q-values). Positions indicate amino acid coordinates within the frameshift-derived sequence. **c.** Distribution of RNA-seq reads overlapping the predicted *zld* frameshift region. Reads were classified according to their alignment to the reference sequence as containing no mismatches, mismatches consistent with sequencing errors, or insertions/deletions.

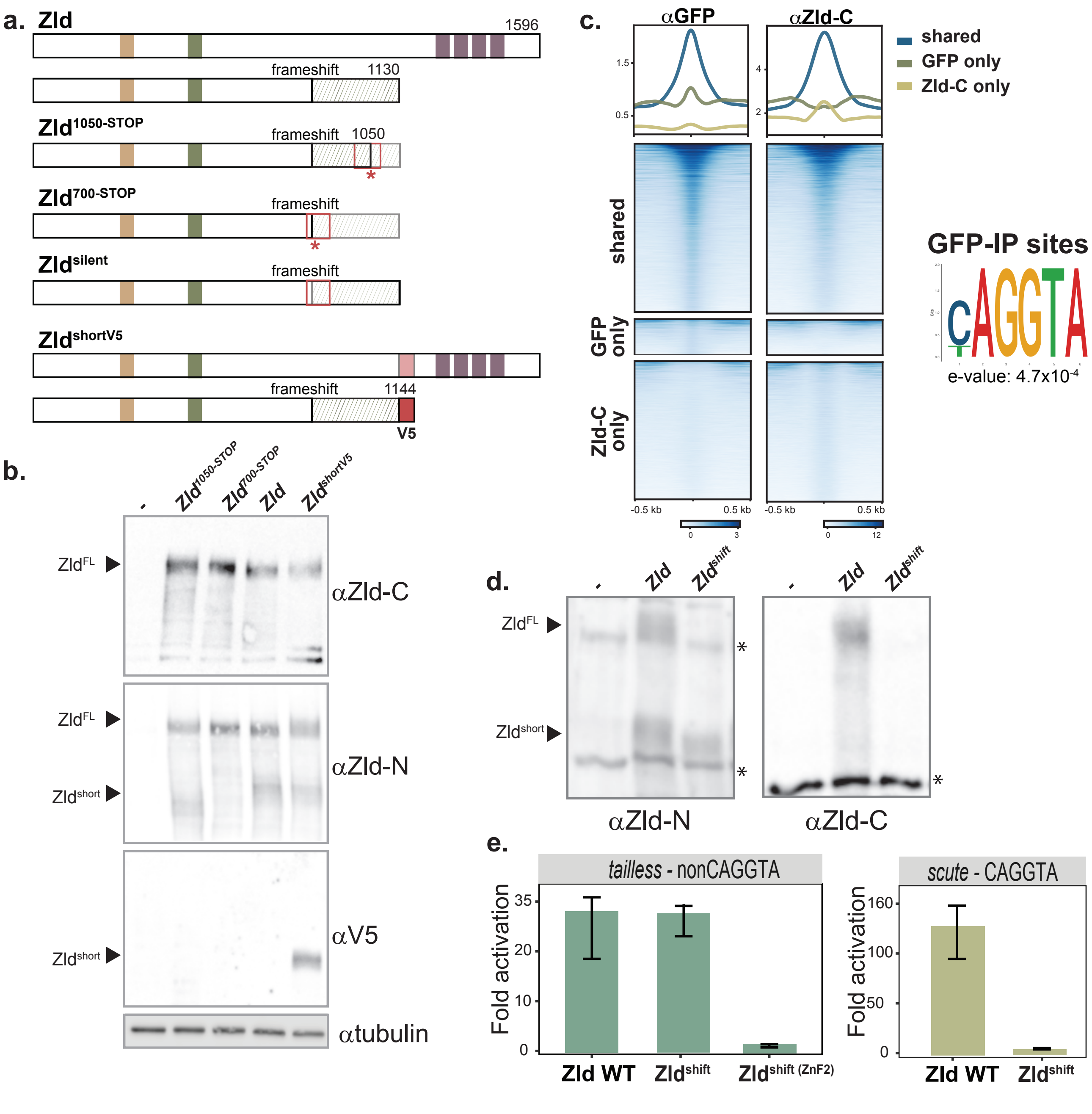


**Extended Data Fig. 8: Forced frameshift produces a functional truncated Zld proteoform. a.** Models of the long and short products that are predicted based on the transfected plasmid in *Drosophila* S2 cells. **b.** Immunoblot for Zld with antibodies against the C-terminus (Zld-C) or N-terminus (Zld-N), V5 or tubulin. Full-length and short products are indicated on the left. **c.** Metaplots of spike-in normalized anti-GFP ChIP-seq. GFP-Zld occupies regions overlapping with those recognized by anti-Zld-C ChIP-seq. Motif enriched in anti-GFP ChIP-seq corresponds to canoncial CAGGTA motif (*right*). **d.** Immunoblot for Zld on S2 cells expressing wild-type Zld or framshifted Zld ($Zld^{shift}$) with antibodies against the N-terminus (αZld-N) or C-terminus (αZld-C). Full-length and short products are indicated on the left. *, non-specific reactivity. **e.** Luciferase reporter assays driven by wild-type Zld, $Zld^{shift}$ or $Zld^{shift}$ with a mutation in zinc finger 2 ($Zld^{shift\ (ZnF2)}$).
